# Deploying dengue-suppressing *Wolbachia: *robust models predict slow but effective spatial spread in *Aedes aegypti*

**DOI:** 10.1101/093229

**Authors:** Michael Turelli, Nicholas H. Barton

**Affiliations:** Department of Evolution and Ecology, University of California, Davis, California, United States of America; Institute of Science and Technology, Am Campus 1, A-3400 Klosterneuburg, Austria

**Keywords:** population transformation, population replacement, bistable wave dynamics, disease suppression, Zika, biocontrol

## Abstract

A novel strategy for controlling the spread of arboviral diseases such as dengue, Zika and chikungunya is to transform mosquito populations with virus-suppressing *Wolbachia*. In general, *Wolbachia* transinfected into mosquitoes induce fitness costs through lower viability or fecundity. These maternally inherited bacteria also produce a frequency-dependent advantage for infected females by inducing cytoplasmic incompatibility (CI), which kills the embryos produced by uninfected females mated to infected males. These competing effects, a frequency-dependent advantage and frequency-independent costs, produce bistable *Wolbachia* frequency dynamics. Above a threshold frequency, denoted *p^̂^*, CI drives fitness-decreasing *Wolbachia* transinfections through local populations; but below *p^̂^*, infection frequencies tend to decline to zero. If *p^̂^* is not too high, CI also drives spatial spread once infections become established over sufficiently large areas. We illustrate how simple models provide testable predictions concerning the spatial and temporal dynamics of *Wolbachia* introductions, focusing on rate of spatial spread, the shape of spreading waves, and the conditions for initiating spread from local introductions. First, we consider the robustness of diffusion-based predictions to incorporating two important features of *wMel-Aedes aegypti* biology that may be inconsistent with the diffusion approximations, namely fast local dynamics induced by complete CI (*i.e*., all embryos produced from incompatible crosses die) and long-tailed, non-Gaussian dispersal. With complete CI, our numerical analyses show that long-tailed dispersal changes wave-width predictions only slightly; but it can significantly reduce wave speed relative to the diffusion prediction; it also allows smaller local introductions to initiate spatial spread. Second, we use approximations for *p̂* and dispersal distances to predict the outcome of 2013 releases of *w*Mel-infected *Aedes aegypti* in Cairns, Australia, Third, we describe new data from *Aedes aegypti* populations near Cairns, Australia that demonstrate long-distance dispersal and provide an approximate lower bound on *p^̂^* for *w*Mel in northeastern Australia. Finally, we apply our analyses to produce operational guidelines for efficient transformation of vector populations over large areas. We demonstrate that even very slow spatial spread, on the order of 10-20 m/month (as predicted), can produce area-wide population transformation within a few years following initial releases covering about 20-30% of the target area.

## 1. Introduction

*Wolbachia* are maternally inherited endosymbionts, pervasive among arthropods (Weinert et al. 2015) and best known for reproductive manipulation (Werren et al. 2008). Their most widely documented reproductive manipulation is cytoplasmic incompatibility (CI) (Hoffmann and Turelli 1997; Hamm et al. 2014), which kills embryos produced by Wolbachia-uninfected females mated to infected males. *Wolbachia-infected* females are compatible with both infected and uninfected males and generally produce only infected progeny. CI gives infected females a reproductive advantage that increases with the infection frequency. Consequently, CI-inducing *Wolbachia* can spread within and among populations, at least once they become sufficiently common that the CI-induced advantage overcomes any frequency-independent disadvantages (Caspari and Watson 1959; Turelli and Hoffmann 1991; Turelli 2010; Barton and Turelli 2011). Because *Wolbachia* are maternally transmitted, selection favors variants that increase the fitness of infected females (Turelli 1994; Haywood and Turelli 2009). Teixeira et al. (2008) and Hedges et al. (2008) discovered that *Wolbachia-infected* individuals are protected from some pathogens, including viruses. Pathogen protection is not universal (Osborne et al. 2009), and studies of both transient somatic *Wolbachia* transinfections (Dodson et al. 2014) and stable transinfections (Martinez et al. 2014) suggest that *Wolbachia* can occasionally enhance susceptibility to pathogens. However, virus protection seems to be a common property of both natural and introduced *Wolbachia* infections (Martinez et al. 2014).

This anti-pathogen effect has revitalized efforts to use *Wolbachia* for disease control, an idea first proposed in the 1960s (Laven 1967; McGraw & O^**’**^Neill 2013). The disease-vector mosquito *Aedes aegypti* has been transinfected with two *Wolbachia* strains from *Drosophila melanogaster* (*w*MelPop,McMeniman et al. 2009; and *w*Mel,Walker *et al*. 2011). Two isolated natural Australian *Ae. aegypti* populations have been transformed with *w*Mel to suppress dengue virus transmission (Hoffmann et al. 2011), and these populations have remained stably transformed for more than four years (Hoffmann et al. 2014; S. L. O **^'^** Neill, pers. comm.). The dengue-suppressing phenotype of *w*Mel-transinfected *Ae. aegypti*, first demonstrated in laboratory colonies (Walker et al. 2011), has been maintained, and possibly enhanced, after two years in nature (Frentiu et al. 2014). Recently, *w*Mel has also been shown to block the spread of the Zika virus by *Ae. aegypti* (Dutra et al. 2016). *Anopheles stephensi* was also transinfected with *Wolbachia*, making them less able to transmit the malaria-causing parasite (Bian et al. 2013). *Wolbachia* transinfections are now being deployed for disease control in at least five countries (Australia, Vietnam, Indonesia, Brazil and Colombia, see the ^“^Eliminate Dengue^”^website: http://www.eliminatedengue.com/program), with many more releases planned. We present simple approximation-based predictions to understand and aid the deployment of these transinfections.

Our mathematical analyses rest on bistable frequency dynamics for *WolbaChia* transinfections. Namely, the frequency-independent costs associated with introduced infections cause frequencies to decline when the infections are rare; but the frequency-dependent advantage associated with CI overcomes these costs when the infections become sufficiently common. As explained in the Discussion, bistability now seems implausible for naturally occurring *WolbaChia* infections (cf.Fenton et al. 2011; Kriesner et al. 2013; Hamm et al. 2014). However, we present several lines of evidence, including new field data, indicating that *w*Mel transinfections in *Ae. aegypti* experience bistable dynamics in nature. Why does bistability matter? As reviewed in Barton and Turelli (2011), bistability constrains which variants can spread spatially, how fast they spread, how difficult it is to initiate spread, and how easily spread is stopped. Roughly speaking, spatial spread can occur only if the critical frequency, denoted *p^̂^*, above which local dynamics predict deterministic increase rather than decrease, is less than a threshold value near ½. As discussed in Turelli (2010), *p^̂^* is determined by a balance between the frequency-dependent advantage provided by cytoplasmic incompatibility and frequency-dependent disadvantages associated with possible deleterious *Wolbachia* effects on fecundity, viability and development time. As *p^̂^* increases, the rate of predicted spatial spread slows to zero (then reverses direction), the area in which the variant must be introduced to initiate spread approaches infinity, and smaller spatial heterogeneities suffice to halt spread. Spatial dynamics depend on details of local frequency dynamics and dispersal that are not well understood empirically. This motivates our exploration of quantitative predictions using relatively simple but robust models that focus on three key biological phenomena, dispersal, deleterious fitness effects and cytoplasmic incompatibility. We seek conditions under which minimal releases of dengue-suppressing *Wolbachia* transinfections achieve area-wide disease control by transforming a significant fraction of the vector population in a relatively short period. We focus on simple models to provide quantitative predictions and guidelines, and test the robustness of the predictions to long-distance dispersal. Our simple approximations make testable predictions that may be improved as additional data become available. Many parameters in detailed models will be difficult to estimate and are likely to vary in time and space. Our idealization is motivated by the scarce information concerning the ecology of disease vectors such as *Ae. aegypti.* For instance, the dynamics of introductions must depend on ecological factors such as density regulation (Hancock et al. 2011a,b). However, the ecology of *Ae. aegypti* is so poorly understood that increases in embryo lethality associated with CI might lead to either decreasing or increasing adult numbers (cf.Prout 1980; Walsh et al. 2013; but see Hancock et al. 2016). As in Barton and Turelli (2011), we ignore these ecological complications and emphasize quantitative conclusions that depend on only two key parameters: σ, a measure of average dispersal distance, and *p^̂^*, the unstable equilibrium frequency. We illustrate how these two parameters can be estimated from introduced*-Wolbachia* frequency data (producing predictions that can be cross-validated) and explore the robustness of the resulting predictions. Our new analyses build on Barton and Turelli (2011), which used diffusion approximations to understand spatial and temporal dynamics. To determine the robustness of those diffusion-based predictions, which make mathematical assumptions that may not be consistent with the biology of *Wolbachia*-transinfected mosquitoes, we examine dispersal patterns that assign higher probabilities to long-distance (and very short-distance) dispersal. We ask how dispersal patterns affect wave speed, wave shape, and the conditions for initiating an expanding wave (Section 4). We use these new, more robust predictions to propose guidelines for field deployment of dengue-suppressing *Wolbachia*. This involves addressing new questions. For instance, Barton and Turelli (2011) determined the minimum area over which *Wolbachia* must be introduced to initiate spatial spread, but ignored the fact that near this critical size threshold, dynamics would be extremely slow. Effective field deployment requires initiating multiple waves to cover a broad area relatively quickly, given constraints on how many mosquitoes can be released. This requires understanding how transient dynamics depend on initial conditions. We synthesize data-based and model-based analyses of spatial spread to outline efficient strategies for area-wide transformation of vector populations (Section 7).

In addition to our theoretical results concerning predicted properties of spatial spread and near-optimal release strategies, we illustrate the theory with predictions concerning the outcome of *w*Mel releases in Cairns, Australia in 2013 (Section 5). We also analyze some previously unpublished data from the 2011 releases reported in Hoffmann et al. (2011) to approximate a lower bound for *p^̂^* relevant to the Cairns releases (Section 6).

## 2. Mathematical background, models and methods

Our initial numerical analyses focus on testing the robustness of predictions presented in Barton and Turelli (2011). We first describe the diffusion approximations and results from Barton and Turelli (2011) before describing the alternative approximations and analyses. Next we describe the model used to analyze the new data we present. Finally we describe our approaches to approximating optimal release strategies.

### 2.1. Diffusion approximations, alternative dynamics and predictions

The simplest spatial model relevant to understanding *Wolbachia* frequency dynamics in space and time is a one-dimensional diffusion approximation:

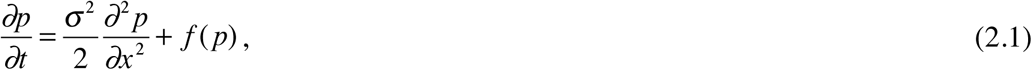

where *f(p)* describes local dynamics and *p(x*,*t*) denotes the infection frequency at point x and time t. If we approximate the local bistable dynamics by the cubic

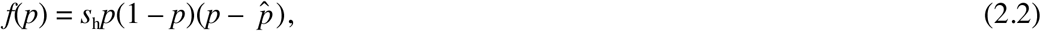

Where *s_h_* describes the intensity of CI, there is an explicit asymptotic traveling wave solution of (2.1), given as Eq. 13 of Barton and Turelli (2011). Eq. (2.1), extended to two dimensions as described by Eq. 22 of Barton and Turelli (2011), can be solved numerically to address transient dynamics associated with local releases. In two dimensions, we interpret *σ*^2^ as the variance in dispersal distance per generation along any axis. (This implies that the average Euclidean distance between the birthplaces of mothers and daughters is 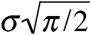assuming Gaussian dispersal.) The model defined by (2.1) and (2.3) provides analytical predictions for wave speed and wave shape and numerical conditions for establishing a spreading wave from a local introduction.

To more accurately approximate CI dynamics, Barton and Turelli (2011) replaced the cubic approximation (2.2) with the model of Schraiber et al. (2012)

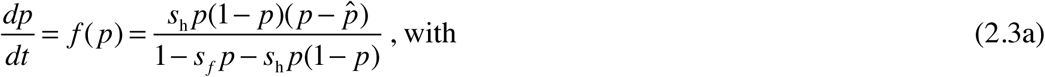

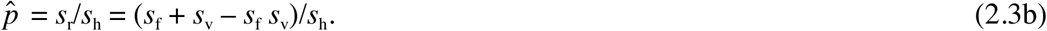

(Eq. 2.3 a assumes that the daily death rate for the infected individuals is *d_I_* = 1, so that time is measured in terms of the average lifetime of an infected individual.) As in Eq. (2.2),*s*_h_ measures the intensity of CI; where as *s_r_* measures the net reduction in fitness caused by the *Wolbachia* infection. As in the discrete-time model of Turelli (2010), fitness costs may involve reductions of both fecundity and mean life length (viability), as measured by *s_f_* and *s_v_*, respectively; however, *s_v_* enters the dynamics only through *p^̂^*. Numerical integration can be used to compare the cubic-based analytical predictions with those produced by this more biologically explicit approximation. For fixed *p^̂^*, the dynamics described by Eq. (2.3a) depend on whether fitness costs primarily involve viability or fecundity effects (because only *s_f_* appears in the denominator). The data of Hoffmann et al. (2014) suggest that fecundity effects may dominate.

In our discrete-time, discrete-space analyses, we approximate local dynamics with the Caspari-Watson model (1959) which incorporates CI and fecundity effects (cf. Hoffmann and Turelli 1988; Weeks et al. 2007). Let *H* denote the relative hatch rate of embryos produced from an incompatible cross. Setting *H* = 1 - *s_h_* and *F* = 1 - *s_f_*, and letting *p* denote the frequency of infected adults, the local dynamics are described by

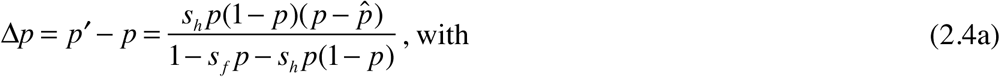

In this model, the condition for bistability (i.e., simultaneous local stability of *p* = 0 and *p* = 1) is > *s_f_* > 0, i.e., the (frequency dependent) benefit to the infection from CI must exceed its (frequency independent) cost, modeled as decreased fecundity. Both lab and field experiments indicate that w Mel causes complete CI, i.e., *s_h_* ≈ 1 in *Ae. aegypti* (Hoffmann et al. 2014).

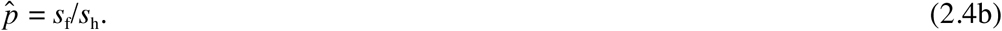

In this model, the condition for bistability (i.e., simultaneous local stability of *p* = 0 and *p* = 1) is > *s_f_* > 0, i.e., the (frequency dependent) benefit to the infection from CI must exceed its (frequency independent) cost, modeled as decreased fecundity. Both lab and field experiments indicate that w Mel causes complete CI, i.e., *s_h_* ≈ 1 in *Ae. aegypti* (Hoffmann et al. 2014).

#### 2.1.1. Wave speed

Measuring time in generations, the predicted wave speed from (2.1) with cubic dynamics (2.1) is

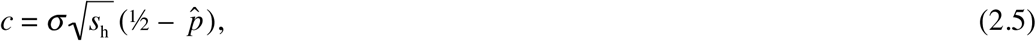

provided that *p* ^**̂**^> 0. This one-dimensional result also describes the asymptotic speed of a radially expanding wave in two dimensions (see Eqs. 23-25 of Barton and Turelli 2011). Barton and Turelli (2011) used numerical solutions of (2.1) to compare this speed prediction to the wave speed produced by (2.3), which explicitly models the fast local dynamics associated with strong CI. The more realistic dynamics (2.3) led to slightly faster wave propagation (as expected because the denominator of *f(p)* is less than one).

#### 2.1.2. Wave width

The explicit traveling-wave solution of (2.1) for cubic f(ρ) provides a simple description for the asymptotic wave width, the spatial scale over which infection frequencies change. Defining wave width as the inverse of the maximum slope of infection frequencies (Endler 1977), the diffusion approximation with cubic dynamics implies that the traveling wave has width

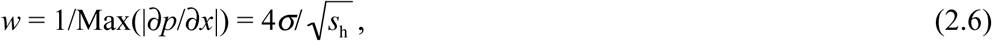

which becomes 4*σ* with complete CI, as in *Ae. aegypti.* The explicit solution that produces (2.6) implies that with *s_h_* = 1, the scaled wave (with space measured in units of *σ)* has shape (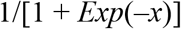) Thus, infection frequencies increase from about 0.18 to 0.82 over 3*σ* If steady spread is observed in the field, we can use this wave-shape approximation to estimate *σ* from spatial infection-frequency data. These estimates can be compared to independent estimates from release-recapture experiments or genetic data. We show below that relation (2.6) is relatively robust to more realistic descriptions of local frequency dynamics and long-tailed dispersal.

#### 2.1.3. Wave initiation

Finally, the diffusion approximation predicts the minimum area that must be actively transformed to initiate deterministic spatial spread. Barton and Turelli (2011) consider introductions over a circular area with initial infection frequency *p_0_* in the circle. This initial state corresponds to rapid local establishment of a transinfection from intensive releases. Hoffmann et al. (2011) showed that releases in isolated suburbs near Cairns produced *w*Mel frequencies over 80% within 12 weeks, about three *Ae. aegypti* generations. Fig. 3 of Barton and Turelli (2011) summarizes the diffusion predictions concerning the minimum radius of release areas, measured in units of dispersal distance *σ*, needed to initiate a spreading wave. In their analysis, the scaled critical radius, denoted. Rent, depends only on *p*^**̂**^. As *p*^**̂**^ increases from 0 to 0.3, *R_crit_* increases from 0 to about 3.5*σ*, then rapidly increases towards infinity as *p* approaches 0.5, the approximate upper bound on *p*^**̂**^ consistent with spatial spread. Releases over areas smaller than R_crit_ are predicted to fail, with the infection locally eliminated by immigration from surrounding uninfected populations. Barton and Turelli (2011) used the Schraiber et al. (2012) model (2.3) to assess the robustness of these cubic-based predictions to more realistic local CI dynamics. Model produced *R_crit_* predictions within a few percent of those derived from the cubic (see Fig. 3 of Barton and Turelli 2011), assuming that *Wolbachia* reduce fitness primarily through viability effects. Below, we contrast the diffusion predictions of Barton and Turelli (2011) with numerical results that account for fecundity effects, faster local dynamics and alternative forms of dispersal.

#### 2.1.4. Time scale for asymptotic wave speed and width

Predictions (2.5) and (2.6) for wave speed and wave width are based on the asymptotic behavior of the traveling wave solutions to the diffusion model (2.1) assuming cubic dynamics. (As discussed in Barton and Turelli (2011), the asymptotic wave speed and width are the same in one dimension and two.) To apply these predictions to frequency data generated from field releases, it is important to know how quickly these asymptotic values are approached. Fig. 1 illustrates numerical solutions for the transient dynamics of the cubic-diffusion model in two dimensions with plausible parameter values for *w*Mel in *Aedes aegypti, s_h_* = 1 and *p*^**̂**^ = 0.25 (discussed below). The calculations use two initial conditions. In the first, the infection is introduced with *p*_0_ near 0.8 over a circular region with diameter 3 (see Fig. 1 legend for details), which is about 14% larger than *R_crit_* = 2.64, the critical radius needed to initiate spatial spread. In the second, the infection frequency drops smoothly from 0.65 at the center of the introduction to 0.25 (the unstable point, *p*^**̂**^) at about *R* = 4.6. As shown in Fig. 1, for these parameters and initial conditions, the approach to the asymptotic wave speed and width is rapid, on the order of five-to-ten generations. Similar results hold for our discrete-time models and data from field releases of *w*Mel in *Aedes aegypti* (data not shown).

**Fig. 1.**
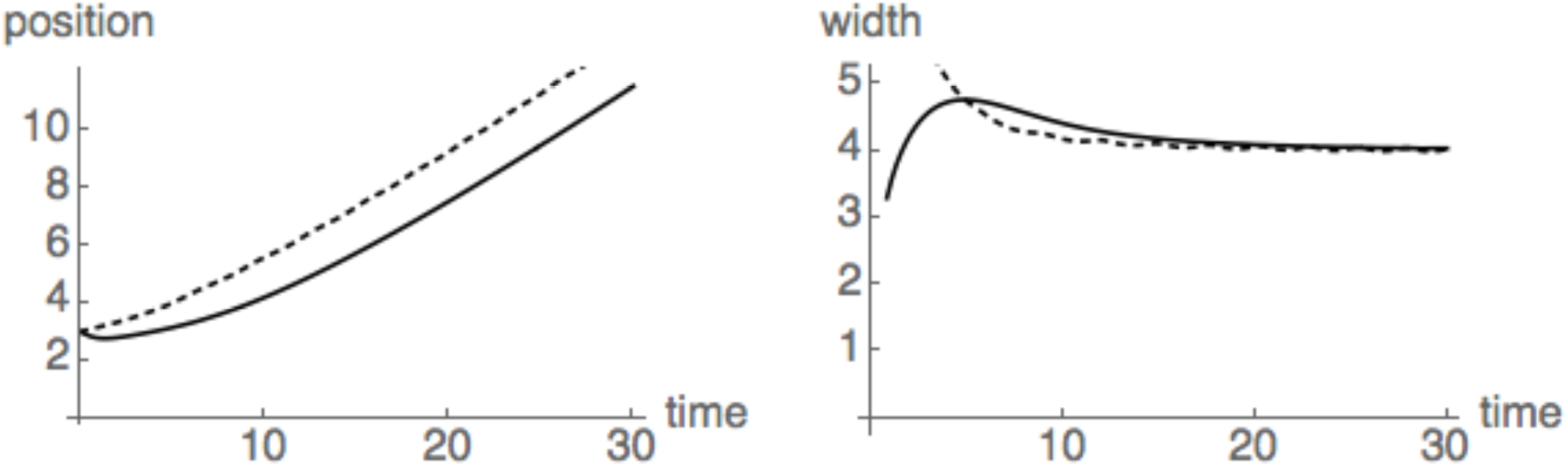
Transient dynamics of wave position and wave width from numerical solutions of the two-dimension version of diffusion model (2.1) with cubic dynamics (2.2), *s_h_* = 1 and *p^̂^* = 0.25. The calculations assume circular introductions with radial symmetry and initial frequency 0.8/{1+exp[4(*R*−3)/*v*]{ for *V* = 0.8 and 8. Setting *V* = 0.8 produces an abrupt drop in the initial frequency from 0.75 to 0.05 over roughly *R* = 2.5 to *R* = 3.5; with *V* = 8, the initial frequency drops from 0.65 at *R* = 0 to 0.25 at *R* = 4.6. The left panel shows wave position measured as the point of maximum slope, the right panel shows the width, measured as the inverse of the maximum slope (see Eq. (2.6)). The dotted curves correspond to *V* = 0.8, modeling a rapid introduction in a confined area. This produces a faster approach to the expected asymptotic speed of 0.25. Both initial conditions, one with narrower width than the asymptotic value of four, the other wider, approach the asymptotic width of four within 7-10 generations.

### 2.2. Numerical analyses of discrete-time, discrete-state (DTDS) models

To explore the robustness of the diffusion predictions, we consider the simultaneous effects of fast local dynamics, associated with complete CI, and long-tailed dispersal. To do this, we replace the PDE approximation (2.1) with discrete-time, discrete-space (DTDS) models that assume discrete generations and discrete patches in which the consequences of mating, fecundity effects and CI occur and between which adult migration occurs.

#### 2.2.1. Model structure and dynamics

Let *i* denote a patch in one or two dimensions, let *g(p) = p*^**'**^ denote a function that describe how mating, fecundity differences and CI transform local infection frequencies between generations, and let *m(i, J)* denote the probability that an individual at location *i* after migration originated in location J. Assuming discrete generations in which migration of newly eclosed individuals precedes local CI dynamics, the infection frequencies among adults in each patch follow

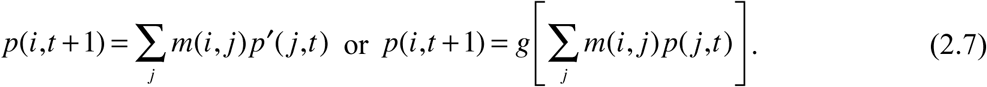

Our choice of patch spacing for these discretizations is discussed below. We approximate local dynamics with the Caspari-Watson model (2.4).

#### 2.2.2. Alternative dispersal kernels

Following Wang et al. (2002), we compare results obtained with three models of dispersal: Gaussian, denoted *G(x)*, versus two ^**“**^long tailed^**”**^ distributions, the Laplace (or reflected exponential), denoted *L(x)*, and the exponential square root (ExpSqrt), denoted *S(x)*. Letting *σ* denote the standard deviation of dispersal distances, our dispersal kernels in one dimension (proportional to the probability of moving distance *x* in one dimension) are

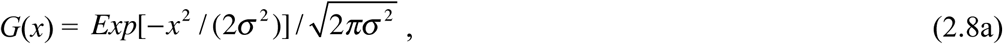

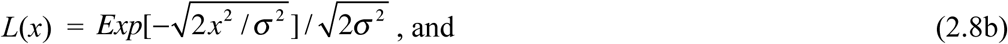

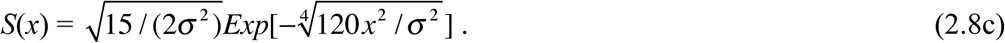

These alternative dispersal models are illustrated in Fig. 2. Our DTDS calculations used patch spacing of 0.5*σ* We truncated the dispersal functions at ±10*σ* We adjusted the variance parameter in our discrete calculations so that the actual standard deviation of the discrete distribution was *σ*. In two dimensions, *(x, y)*, the dispersal models, generically denoted *m(z)*, were implemented as 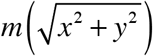

**Fig. 2:**
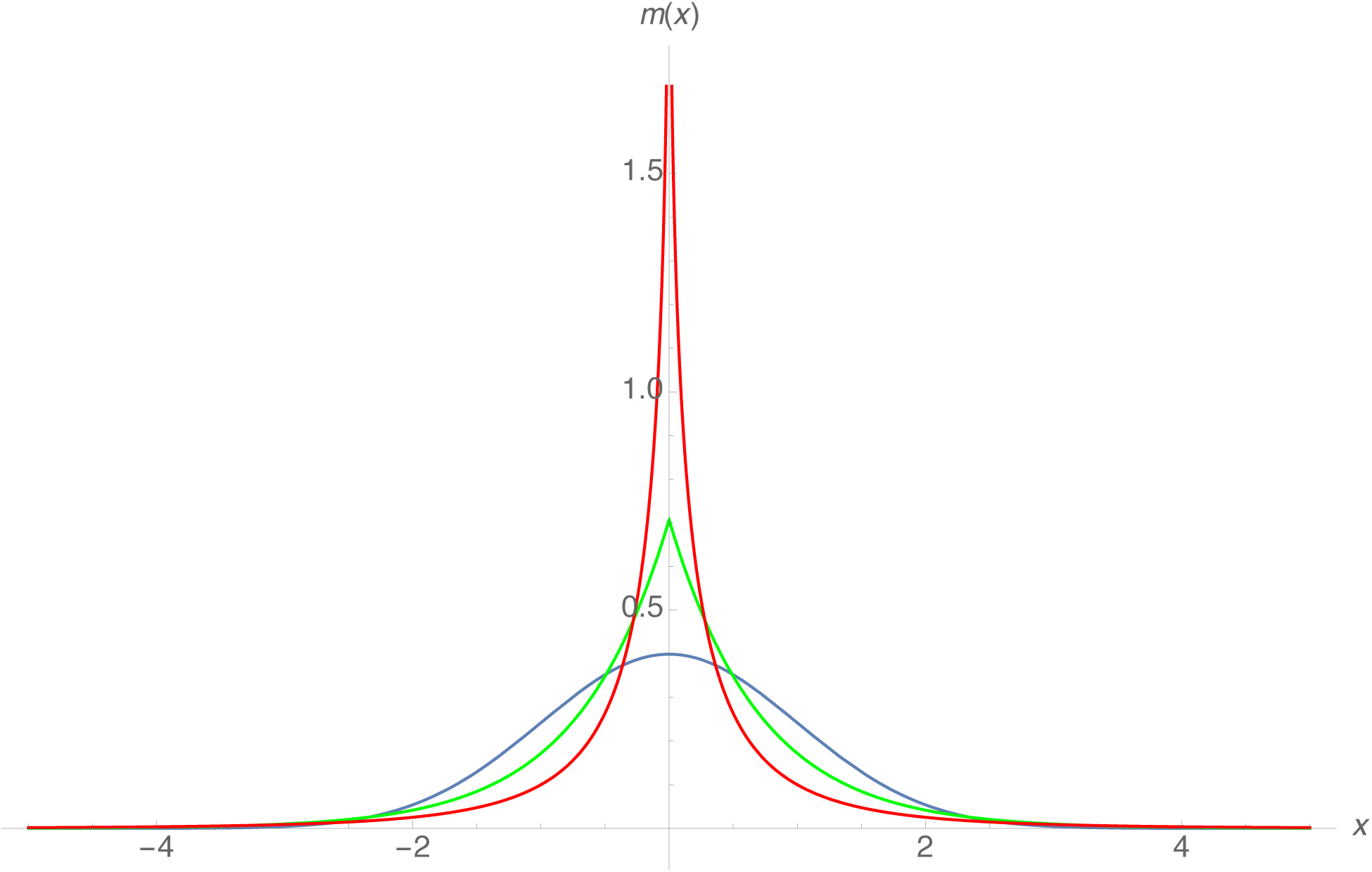
Alternative dispersal models with σ= 1. The three models are: Gaussian (blue), Laplace (green), and ExpSqrt (red) as described by (2.8). Each describes the probability, denoted m(x) in the figure, of moving distance *x* along any axis.

**Fig. 3.**
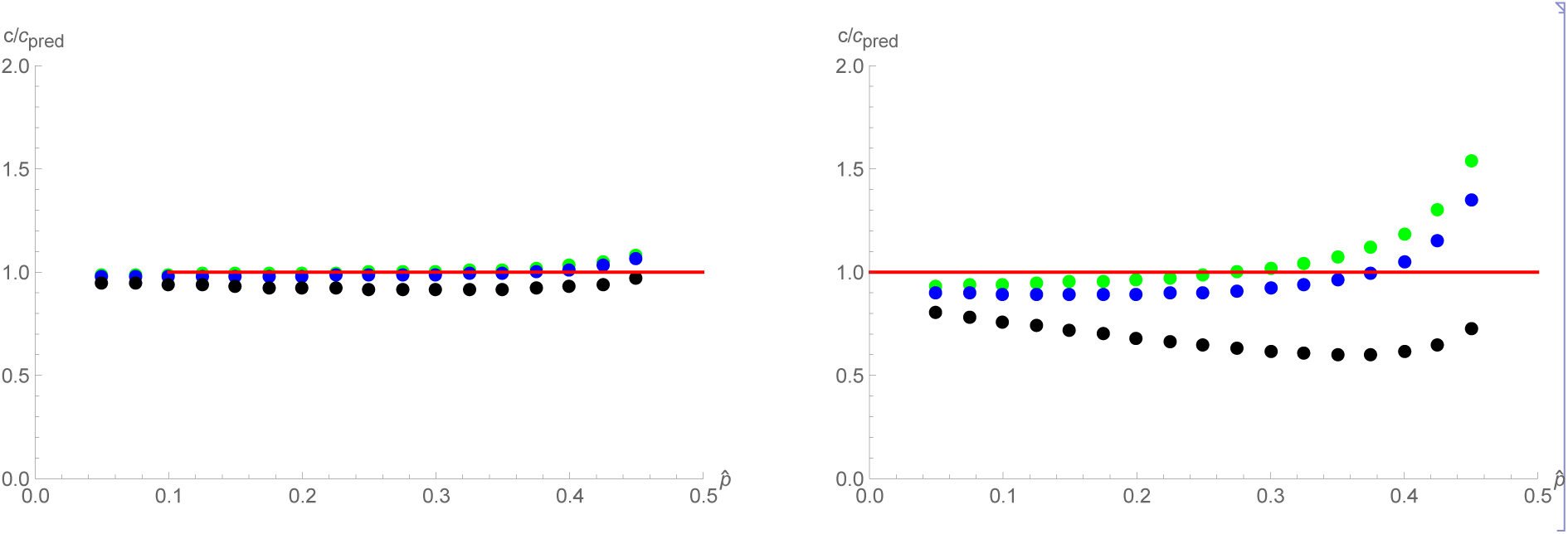
Wave speed. Speed calculated from DSDT analyses compared to the cubic-based diffusion prediction, 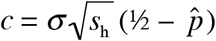 as a function of *p^̂^*, for *s_h_* = 0.2 (left) and *s_h_* = 1 (right).

Taking logs of the densities, the tails of *G(x)* decline as -*x*^2^, whereas the tails of *L(x)* and S(x) decline as -|*x*| and 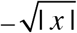, respectively, corresponding to successively higher probabilities of long-distance dispersal. Denoting the random variables corresponding to these densities as G, *L* and S, we have *P*(|*G*|> 3*σ*) = 0.003,*P*(|*L*|> 3*σ*) = 0.014 and *P*(|*S*|> 3*σ*)=0.002. As Fig. 2 shows, higher probabilities of long-distance dispersal are accompanied by higher probabilities of short-distance dispersal (e.g.,*P*(|*G*|< 0.5 *σ*)= 0.383,|*L*|> 0.5 *σ*)= 0.507 and *P*(|*S*|< 0.5 *σ*)= 0.678), with corresponding medians for |*G*|, |*L*| and |*S*| of 0.67*σ*, 0.49σand 0.28*σ*, respectively.

In two dimensions, we assume that dispersal is isotropic, with radial distribution given by the kernels defined by (2.8), and scaled such that the standard deviation along any one axis is *σ*. To approximate *σ* from experiments that estimate mean Euclidean dispersal distances, D, note that the Gaussian produces 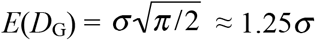whereas for the Laplace and ExpSqrt, we have E(D_L_) ≈ 1.15*σ* and E(D_s_)≈ 0.94σ, respectively. Thus, empirical estimates of average Euclidean dispersal distance can imply values of *σ* that differ by over 30% depending on the shape of the dispersal function, with longer tails implying higher values of *σ*. (Note that statistical estimation of dispersal requires some assumption about the distribution of dispersal distance.) We compare the predictions resulting from alternative dispersal models by holding fixed the variance parameter σ^2^, which is natural measure of dispersal distance for diffusion approximations (see, for instance, the derivations in Haldane (1948), Slatkin (1973) or Nagylaki (1975)).

### 2.3 Model used for data analysis: an isolated deme subject to immigration

Barton and Turelli (2011) adapted the ^**“**^island model^**”**^ of Haldane (1930) to approximate the rate of immigration of *Wolbachia-infected* individuals required to ^**“**^flip^**”**^ an isolated population from uninfected to infected. In addition to approximating the critical migration rate, *m*, the analysis produces an analytical approximation for the equilibrium infection frequency when the immigration rate is too low to flip the recipient population to *Wolbachia* fixation. Assuming complete CI, 100% frequency of *Wolbachia* in the donor population and one-way immigration into the recipient population, Eq. (31) of Barton and Turelli (2011) predicts that *Wolbachia* should take over the recipient population if m, the fraction of individuals who were new migrants each generation, exceeds m* = p^̂2^ / 4. For *m* < m*, the predicted *Wolbachia* equilibrium frequency in the recipient population, using a cubic approximation for local dynamics, is

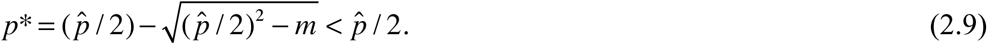

(this is a reparameterization of Eq. (31) of Barton and Turelli 2011). Hence, if we use the longterm average *Wolbachia* frequency, *p─*, to approximate p*, the equilibrium described by (2.9), we can approximate a lower bound for the unstable equilibrium, *p*^̂^, as 2 *p─*.

### 2.4. Near-optimal release strategies

We analyze alternative release strategies using a combination of numerical solutions of diffusion models, DTDS models and even simpler models that assume constant rates of radial spread from release foci. Each analysis is described below along with the results it produces.

## 3. New data demonstrating bistability

We analyze a small subset of the *Wolbachia* infection frequency data collected subsequent to the first ^**“**^Eliminate Dengue^**”**^ field releases of *w* Mel-infected *Ae. aegypti*, described in

Hoffmann et al. (2011). The releases occurred in early 2011 in two isolated towns, Gordonvale (668 houses) and Yorkeys Knob (614 houses), near Cairns in northeast Australia. As described in Hoffmann et al. (2011), Pyramid Estate (PE) is an area of Gordonvale separated from the town center by a major highway, with roughly 100 m separating the nearest houses on either side. Highways seem to inhibit *Ae. aegypti* migration (Hemme et al. 2010). The 2011 *w*Mel releases were restricted to the main part of Gordonvale; but as reported in Hoffmann et al. (2011), *w*Mel-infected mosquitoes were found in PE within months of the initial releases. The PE capture sites were scattered over an area of houses on the order of 1 km^2^ with traps roughly 100-500 m from the nearest residences in the Gordonvale release area. As described in Hoffmann et al. (2011,2014), infection frequencies were estimated using PCR of DNA from adults reared from eggs collected in oviposition traps. Between late March 2011 and January 2015, 2689 adults were assayed in PE. The data are archived in Dryad (http://XXX).

## 4. Results: Robustness of diffusion results to long-tailed dispersal and rapid CI dynamics

### 4.1. Wave speed

We initially calculated wave speed in a one-dimensional spatial array, then as in Barton and Turelli (2011), we checked the results with two-dimensional calculations. To disentangle the effects of non-Gaussian dispersal from the effects of fast local dynamics, we contrast results obtained assuming complete CI (*s_h_* = 1), as observed with *w*Mel-infected *Ae. aegypti*, with results assuming weak CI (*s_h_* = 0.2). Fig. 3 compares the numerically approximated wave speeds to the analytical prediction, 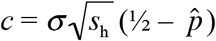 from the diffusion approximation with cubic dynamics. The left panel shows that with relatively slow local dynamics (*s_h_* = 0.2), the cubic diffusion approximation is accurate and robust to the shape of the dispersal function. This is expected from the derivation of approximation (2.1) as a limit of discrete-time, discrete-space dynamics (Haldane 1948; Nagylaki 1975). The derivation explicitly invokes slow local dynamics and limited dispersal, retaining only the variance of dispersal distances in the quadratic approximation. The s_h_= 0.2 results have two other notable features. First, despite the overall accuracy of approximation (2.5), we see that ExpSqrt dispersal slightly slows propagation. This can be understood in terms of the lower median dispersal distance and the fact that with bistability, rare long-distance dispersal is not effective at pushing the wave forward, because long-distance migrants are swamped by the much more abundant natives. This distinguishes bistable spatial dynamics from those with zero as an unstable equilibrium. For such systems, long-tailed dispersal can produce accelerating waves (see Supporting Information Appendix A for references and comparison of bistable versus Fisherian wave speeds). Moreover, geographic spread associated with spatially non-contiguous, successful long-distance colonization events (cf.Shigesada and Kawasaki 1997, Ch. 5), can greatly exceed predictions based on average dispersal distances. Second, note that as *p^̂^* approaches 0.5, the analytical approximation starts to underestimate wave speed. As described by Barton and Turelli (2011), this reflects the fact that the cubic model produces the threshold *p^̂^* < 0.5 for spatial spread, whereas models more accurately describing CI and fitness costs, such as (2.3) and (2.4), predict spatial spread with *p^〠^*slightly above 0.5.

The green dots were produced with Gaussian dispersal, blue with Laplace and black with Exponential Square root (ExpSqrt).

The right panel of Fig. 3 *(s_h_* = 1) shows that faster local dynamics accentuate both phenomena seen with = 0.2: slower speed with more long-tailed dispersal and underestimation of observed speed as *p^̂^*;approaches 0.5. With complete CI and plausible *p̂* (*i.e.*,0.2≤ *p*̂≤ 0.35), observed speed closely follows the cubic-based diffusion prediction with Gaussian dispersal, but is reduced by about 10% for Laplace dispersal and by much more (25-40%) for ExpSqrt dispersal. A simple interpretation is that long-tailed dispersal is associated with smaller median dispersal distances. Long-distance migrants are effectively ^**“**^wasted^**”**^ in that they cannot initiate local spread.

Appendix A provides a more complete description of the consequences of alternative dispersal models on wave speed under bistable versus monostable local dynamics, including the consequences of finite population size at the front on wave propagation.

### 4.2. Wave width

Under the diffusion model with cubic dynamics, the predicted wave width is 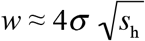 (2.6). Fig. 4 compares this prediction with the results obtained from DTDS with Caspari-Watson dynamics and alternative dispersal models. With relatively slow local dynamics (*s_h_* = 0.2), Panel A shows that the cubic-diffusion prediction remains accurate for all three dispersal models, analogous to the results for wave speed illustrated in Fig. 3A. ExpSqrt dispersal slightly reduces wave width, presumably reflecting the lower median dispersal. As with wave speed,*s_h_* = 1 produces larger departures from the cubic-diffusion prediction and much greater effects of dispersal shape. However, for plausible values of *p^̂^* (i.e, 0.2 < *p^̂^*; < 0.35), the observed width remains within about 15% of prediction (2.6) for Gaussian and Laplace and very close to the prediction for ExpSqrt.

**Fig. 4.**
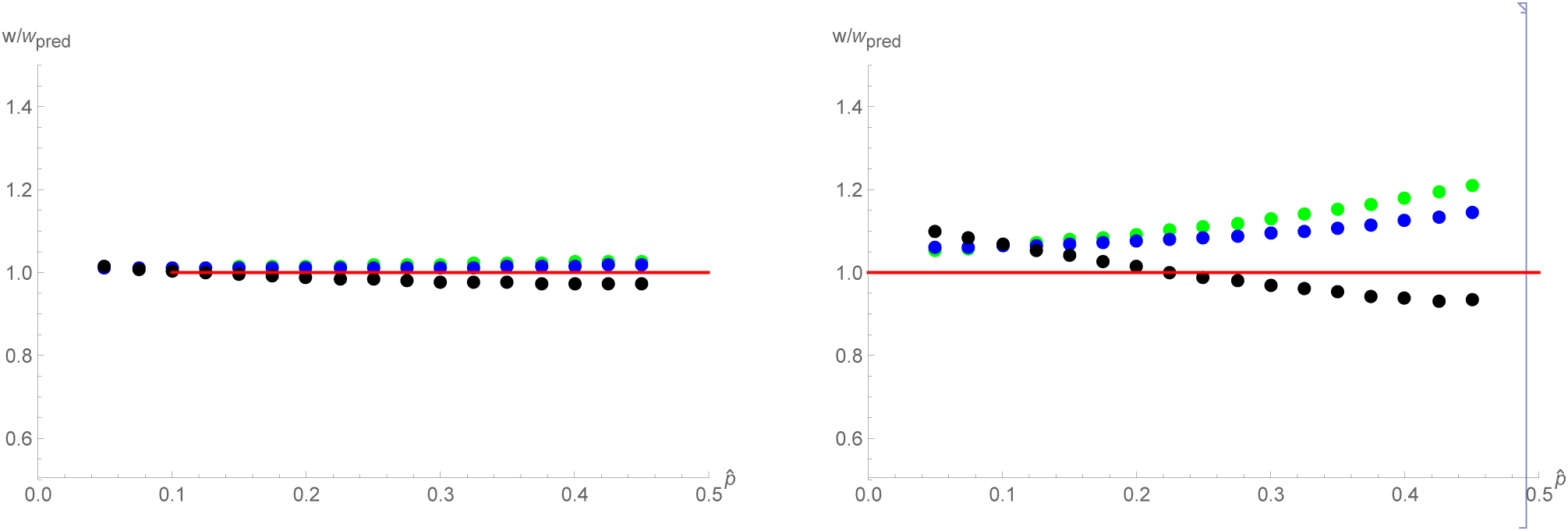
Wave width. Wave width as a function of *p* calculated using discrete-time, discrete-space (DTDS) analyses with alternative dispersal models. The dots from the DTDS analyses are compared to the diffusion prediction 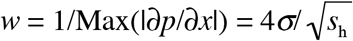 (right). The green dots were produced with Gaussian dispersal, blue with Laplace and black with Exponential Square root (ExpSqrt).

### 4.3. Wave initiation: critical radius R_cr_it

Fig. 3 of Barton and Turelli (2011) showed how R_cr_it, the minimal radius of an introduction needed to initiate spread (measured in units of the dispersal parameter σ), depends on *p0* and *p_0_* under the diffusion approximation. It contrasts the predictions for cubic dynamics versus Schraiber et al. (2012)*Wolbachia* dynamics (Eq. 2.3). Fig. 5 compares those predictions to DTDS results under Caspari-Watson *Wolbachia* dynamics (Eq. 2.4). The key result is that longtailed dispersal produces smaller critical radii, and the effect of long-tailed dispersal increases as *p* increases. This result is complementary to the wave-speed results. With longer-tailed dispersal, more individuals move very little so that the median dispersal falls, making it easier to establish a wave (but the resulting wave moves more slowly). The discrepancies between the diffusion results with Schraiber et al. (2012) dynamics and the DTDS results for Gaussian dispersal are mainly attributable to the fact that the Schraiber et al. (2012) results illustrated in Fig. 5 assume only viability costs, which produces slower dynamics (see Eq. 2.4 a) and requires larger introductions, than if one assumes fecundity costs, as done in the DTDS Caspari-Watson model. The effect of fecundity vs. viability costs is illustrated in Table 1 in section 5.1. For *p̂* = 0.35 and p_0_ = 0.8, numerical solution of the diffusion equation with Schraiber et al. (2012) dynamics produces *R_crit_* = 3.36 if *s_f_* = (so that *p^̂^* = s_v_), but this drops to R_cri_t = 2.76 if s_v_ = 0 (so that *p^̂^* = s_f_). The corresponding values under the DTDS model with Caspari-Watson dynamics are R_cri_t = 3.01, 2.91, 2.55, for Gaussian, Laplace and ExpSqrt dispersal, respectively.

**Fig. 5.**
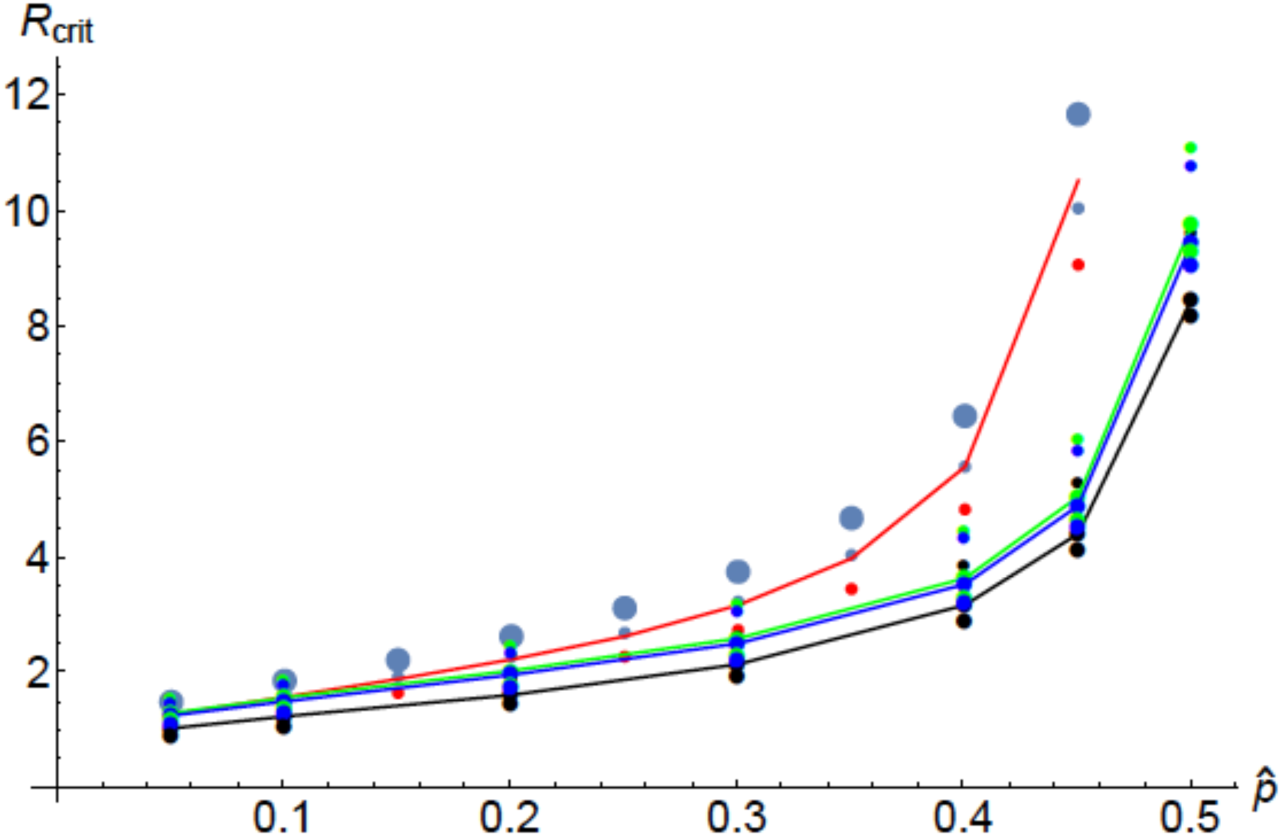
Critical radius, R_crit_, assuming complete CI and alternative dynamics. The upper points reproduce the diffusion results from Barton and Turelli (2011) with cubic (red curve) and Schraiber et al. (2012) CI dynamics (assuming only viability fitness costs) The red curve shows the cubic-diffusion predictions with p_0_ = 0.8, the large blue dots are the cubic with p_0_ = 0.6. The small red (blue) dots are produced by the diffusion analysis of Schraiber et al. (2012) CI dynamics with p_0_ = 0.8 *(p_0_* = 0.6). The lower points and curves show our DTDS predictions as a function of *p* and the initial infection frequency (p_0_) with alternative dispersal models. The lower lines correspond to p_0_ = 0.8 with Gaussian (green), Laplace (blue) and ExpSqrt (black) dispersal. The points above and below these lines correspond to p_0_ = 0.6 and p_0_ = 1, respectively

There are two striking results concerning the DTDS-derived values of R_cri_t. First, like the wave-width results, the critical radii are relatively insensitive to the dispersal model. Second, however, unlike the wave-width results, the critical radii are significantly different and smaller than those produced by the diffusion approximation. Barton and Turelli (2011) showed that the Schraiber et al. (2012) dynamics produced smaller R_crit_ values than the cubic model, even if fitness costs were purely based on reduced viability. Reduced fecundity, as assumed in the Caspari-Watson model, accelerates the local dynamics and hence allows much smaller introductions to initiate a traveling wave. Even with *p^̂^* = 0.35, introductions with *p_0_* = 0.8 will succeed as long as the initial radius of release, denoted R_I_, satisfies R_I_ > 2.5*σ*(or 3.0σ) with ExpSqrt (or Gaussian) dispersal.

## 5. Results: Predictions for 2013 Cairns releases

### 5.1. Diffusion-based predictions

One of our primary aims is to understand the robustness of the Barton and Turelli (2011) diffusion predictions. Rather than discuss generalities, we will focus on specific field releases. In early 2013, three localized releases were performed within the city of Cairns. Releases were made in three neighborhoods, Edgehill/Whitfield (EHW), Parramatta Park (PP), and Westcourt (WC). The release areas were roughly 0.97 km^2^ for EHW, 0.52 km^2^ for PP, and only 0.11 km^2^ for WC. Infection frequencies quickly rose above 0.8 within all three release areas, and each release area adjoined housing into which the *w*Mel infection might plausibly spread. What predictions emerge from the diffusion approximations?

Numerical predictions require estimates of *σ* and *p^̂^*. Russell et al. (2005) performed a mark-release-recapture experiment with *Ae. aegypti* using a release site abutting the 2013 EHW release area. The mean absolute distance of recaptures from the release point was about 78 m. The diffusion approximation assumes that dispersal is measured as the standard deviation of dispersal distance along any axis. If we assume that dispersal distance is roughly Gaussian distributed with mean 0 and standard deviation *σ* along each axis, the mean absolute dispersal distance is 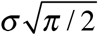 or about 1.25σ. With this assumption, the estimate from Russell et al. (2005) implies *σ* ≈ 62 m/(generation). In general, however, release-recapture estimates tend to be systematically lower than those based on genetic data (see, for instance, Barton and Hewitt 1985, Fig. 3). Moreover, estimates of dispersal distance for *Ae. aegypti* are extremely variable in time and space. For instance, Harrington et al. (2005) found that repeated estimates of mean dispersal distance in the same village in Thailand ranged from about 40 m/(generation) to about 160 m/(generation)^1/2^. Our theoretical predictions concerning the consequences of dispersal are best interpreted as temporal averages, which are more likely to be accurately captured by indirect estimates of average dispersal such as wave width (or genetic data describing the decline of relatedness with distance). Given that direct estimates systematically underestimate average dispersal in nature, we use *σ≈100* m/(generation) as a plausible estimate for Cairns. We recognize, however, that dispersal is likely to vary significantly with local conditions.

Assuming *σ≈* 100 m/(generation), Eq. (2.6) implies that if spatial spread is observed, the wave width should be about 400 m. From Eq. (2.5), the corresponding wave speed is *C* = 100(1/2 – *p^̂^*) m per generation (m/gen). As argued in section 6.1 below, *p^̂^* is probably above 0.2. Thus, the maximum predicted speed is about 30 m/gen. However, if *p* is as high as 0.35, predicted speed falls to 15 m/gen. Assuming about 10 *Ae. aegypti* generations per year near Cairns, these crude estimates indicate that *w*Mel spread in *Ae. aegypti* is likely to be on the order of 150-300 m/year - two or three orders of magnitude slower than the spread of *w*Ri in California and eastern Australia *D. simulans* (100 km/year,Kriesner et al. 2013). Yet repeated estimates of dispersal distances for various *Drosophila* species suggest that natural dispersal distances are at most 5-10 times greater for *D. simnlans* than for *Ae. aegypti* (*e.g*.,Dobzhansky and Wright 1943; Powell et al. 1976; McInnis et al. 1982). The critical difference between the speeds associated with these exemplars of *Wolbachia* spread is unlikely to be dispersal, but more probably the bistability of *w*Mel dynamics in *Ae. aegypti* versus the monostability of *w*Ri dynamics (see Discussion section 8.3). Monostability allows relatively rare human-mediated, long-distance dispersal to greatly enhance spatial spread, as described, for instance, by ^**“**^structured diffusion^**”**^ models (Shigesada and Kawasaki 1997, Ch. 5).

Whether spatial spread occurs with bistability depends on the size of the release area, the initial frequency produced in the release area (p_0_), and *p^̂^*. From Fig. 3 of Barton and Turelli (2011) withp_0_ = 0.8, if *p* were as large as 0.35, the minimum radius of a circular release needed to produce an expanding wave would be on the order of 4σ, implying a minimal release area of about 0.5 km^2^ (assuming *σ* ≈ 100 *m*/(generation)^1/2^). Replacing the cubic in (2.2) with the Schraiber et al. (2012) description of CI dynamics (2.3), Barton and Turelli (2011) showed that the minimal radius with *p^̂^* = 0.35 falls from about 4*σ* to about 2.8-3.5*σ*, with the value depending on whether *w*Mel-infected *Ae. aegypti* lose fitness primarily through fecundity (as the data of Hoffmann et al. 2014 suggest), which produces 2.8*σ*, or viability, which produces 3.5*σ* The smaller values (from Schraiber et al. 2012) imply minimal release areas of about 0.25-0.38 km^2^ (the lower value assumes only fecundity effects). In contrast, if *p^̂^* were as small as 0.2, the minimum radius falls to about 2*σ* both the cubic model and Schraiber et al. (2012) dynamics (with either fecundity or viability effects), corresponding to a minimal area of about 0.13 km^2^.

Table 1 summarizes our diffusion-based predictions. Note that according to these analyses, the releases at EHW and PP should certainly lead to spatial spread, but the WC release is close to minimal release area even if *p* is as small as 0.2. Next we address the robustness of these predictions to long-tailed dispersal and patchy spatial distributions.

**Table 1.** Diffusion-based predictions for spatial spread.

| Unstable point | Speed (m/gen) | Width (m) | Minimum Release Area (km<sup>2</sup>) |
| --- | --- | --- | --- |
| $\hat{p} = 0.2$ | 30 | 400 | 0.13 |
| $\hat{p} = 0.35$ | 15 | 400 | 0.25-0.38<sup>a</sup> (0.5) |

<sup>a</sup>The smaller prediction (0.25) is derived with Schraiber et al. (2012) CI dynamics (2.3) assuming that wMel reduces only the fecundity of *Ae. aegypti* (i.e., $s_f = s_f$ in 2.3b), the larger result (0.38) assumes that wMel reduces only viability. The cubic model produces the still larger value (0.5).

### 5.2. DTDS-based predictions

Our robustness analyses of the wave-width predictions emerging from the cubic-diffusion model indicate that *σ* can be reliably estimated from observed widths of traveling waves of *Wolbachia* infections. In contrast, our wave-speed analyses suggest that given an estimate of *σ*, the predicted wave speed depends significantly on the shape of dispersal with plausible speeds that may be on the order of 20-30% below the cubic-diffusion prediction 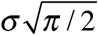

Our final prediction concerns spatial spread from individual localized releases. As shown in Fig 5, the critical release radius for spread depends on: 1) *p^̂^*, the unstable point; 2)p_0_, the initial infection frequency produced within the release areas; 3) the shape of the dispersal function; and 4) *σ*, dispersal distance. As dispersal becomes more long-tailed (moving from Gaussian to ExpSqrt), the critical radius of the initial introduction decreases. If we assume that *p^̂^* = 0.3 and p_0_ = 0.8, R_crit_ is about 2.61 *σ* if dispersal is Gaussian, but falls to about 2.51 *σ* (or 2.16*σ*) if dispersal is Laplace (or ExpSqrt). Hence, for each of the three release areas in Cairns, we can ask what is the maximum *σ* consistent with our deterministic predictions for spatial spread. Given that spread occurs only if the release area exceeds 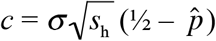 for each release area, we can approximate an upper bound on *σ* consistent with spatial spread by

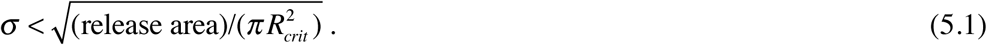

Table 2 presents these upper bounds on *σ* associated with the three 2013 release areas in central Cairns for a plausible range of *p^̂^*

**Table 2.** Predicted maximum σ(in meters) consistent with spatial spread.

| Location | $\hat{p}$ | Gaussian | Laplace | ExpSqrt |
| --- | --- | --- | --- | --- |
| Westcourt (0.11 km<sup>2</sup>) | 0.2 | 92 | 95 | 115 |
|  | 0.25 | 81 | 84 | 101 |
|  | 0.3 | 72 | 74 | 87 |
|  | 0.35 | 62 | 64 | 73 |
| Parramatta Park (0.52 km<sup>2</sup>) | 0.35 | 135 | 140 | 160 |
| Edge Hill/Whitfield (0.97 km<sup>2</sup>) | 0.35 | 184 | 191 | 218 |

These prediction, based on inequality (10), assume that assuming that $p_0 = 0.8$ and $0.2 \leq \hat{p} \leq 0.35$ .

Given that very few empirical estimates of *σ* for *Aedes aegypti* exceed 100 m, these results suggest that spatial spread should certainly be observed for the Edge Hill/Whitfield and Parramatta Park releases. The prediction for Westcourt is more ambiguous. Note that from Table 1, our diffusion predictions with 0.2 < *p^̂^* < 0.35 indicated a minimum release area of 0.14 km^2^. This lower bound assumes *σ* = 100 m and *p^̂^* = 0.2. Thus the diffusion analyses suggested probable failure of the Westcourt release. In contrast, as shown in Fig. 5, our DTDS analyses indicate that the Westcourt release area may be near the lower limit for spread, with the outcome depending critically on the exact values of *σ* and *p^̂^*.

Empirically testing these predictions concerning minimal release areas is confounded by the fact that dynamics very close to the critical values for spread are expected to be slow. Assuming that *p^̂^* = 0.25, if the release area is 10% (5%) smaller than the critical value, the time for collapse is on the order of 15-20 (20-25) generations, roughly two years. Conversely, if the release area is only 10% (5%) larger that the critical area, the time scale for appreciable spatial spread is also on the order of 15-20 (20-25) generations. In contrast, release areas twice as large as necessary should produce appreciable spread in only 10-15 generations; whereas release areas only half as large as needed should essentially collapse in 10-15 generations. These calculations motivated our analyses presented below of ^**“**^optimal^**”**^ release sizes aimed at area-wide coverage within a few years.

## 6. Results: Data relevant to bistability and long-distance dispersal

### 6.1. Heuristic approximation for p^̂^ from Pyramid Estates data

Pyramid Estates (PE) was sampled for over two years after the releases stopped. The few capture sites were scattered over an area of houses that is on the order of 1 km^2^ with traps varying between about 100 m and 500 m from the nearest residences in our release area. For over two years, the *w*Mel frequency in PE remained persistently low, but non-zero with *p*≈ 0.106 (N = 2689, averaged over space and time). (We found no evidence that infection frequency varied with distance from the release area). For instance, a sample of 43 *Ae. aegypti* from the week ending 9 January 2015 yielded an infection frequency of 0.07 [with 95% binomial confidence interval (0.01, 0.19)]. From Eq. (2.9), a long-term average of 0.105 implies *p^̂^* > 0.21. The persistence of a low infection frequency for over two years clearly demonstrates regular immigration of infected individuals that has been unable to push the local PE population past its unstable point. The fitness data from Hoffmann et al. (2014) suggest that *p* for *w*Mel near Cairns is likely to be at least 0.2. This is corroborated by the transient dynamics described in Hoffmann et al. (2011) which also suggest that *p* ^*̂*^ is unlikely to be significantly above 0.3.

### 6.2. Long-tailed dispersal

Gordonvale and Yorkeys Knob are separated from other sizable populations of *Ae. aegypti* by kilometers. Yet, Hoffmann et al. (2014) found consistent low frequencies of uninfected individuals more than three years after *w*Mel reached near-fixation, despite no evidence for imperfect maternal transmission. Yorkeys Knob is less isolated than Gordonvale and shows a significantly higher frequency of uninfected individuals, about 6% versus 3%. Long-distance dispersal is the most plausible explanation for uninfected individuals in Gordonvale and Yorkeys Knob - and the persistence of rare infected individuals at Pyramid Estate.

## 7. Results: Near-optimal release strategies

We seek conditions under which releases of disease-suppressing Wolbachia transinfections achieve area-wide control of a disease such as dengue (cf.Ferguson et al. 2015) by transforming a significant fraction of the vector population, say 80%, in a relatively rapid period, say two to four years (on the order of 20-40 generations), while releasing as few Wolbachia-infected vectors as possible. We consider several questions associated with the optimizing the timing, spacing and intensity of releases. First, we contrast pulsed releases, over a time scale of very few vector generations, with prolonged low-intensity releases. Second, we consider optimizing the spacing and intensity of releases, as quantified by three parameters: a) local initial infection frequencies after releases, b) areas of local releases, and c) the spacing of releases. Third, given that optimization requires knowing parameters that can only be approximated, we consider the consequences of non-optimal releases.

### 7.1. Timing of releases: pulse versus gradual introduction

With bistable dynamics, the frequency of an infection (or allele) must be raised above a critical threshold, *p^̂^*, over a sufficiently large area to initiate spread. What is the most efficient way to establish an infection? At one extreme, the frequency could be raised essentially instantaneously to some p_0_(x); ifp_0_ > *p^̂^* over a large enough region (cf. Fig. 5), the infection will spread. At the other extreme, there might be a gradual introduction, described by a local introduction rate M(X)sustained until deterministic spread is initiated. If this input is sufficiently high over a sufficiently large region, the infection will be locally established and spread.

Between these extremes, releases might be sustained for a set period of many months or a few years. Supporting Information Appendix B investigates conditions for local establishment and wave initiation, providing analytical results for a single deme and for a point source of introduction in one dimension, and numerical results for two dimensions. We show that it is most efficient to raise infection frequency rapidly, in a brief pulse, rather than making gradual introductions. This accords with the intuition that it is most efficient to raise the frequency as quickly as possible above the threshold *p^̂^*: this maximizes the reproductive value of introduced individuals. The principle is simple; during gradual introductions, until local infection frequencies exceed *p^̂^*, the introduced infected individuals are systematically eliminated by deterministic selection that dominates the weaker (frequency-dependent) force of CI at low *Wolbachia* frequencies. Assuming that releases quickly drive the local infection frequency to a value po sufficient to initiate spatial spread, we ask how long it might take to cover a large area and what spatial patterns of release minimize the time to reach a desired coverage.

### 7.2. Spacing and intensity of releases

We start with idealized analyses, then discuss their relative robustness and the effects of environmental heterogeneity. Consider an area with a relatively uniform vector density. What is the optimal release strategy? The calculations in Appendix B show that for a given number of mosquitoes, the best strategy is to release a short pulse, *i.e*, to essentially instantly produce a local infection frequency sufficient to initiate a wave. Obviously there are practical constraints on numbers that can be released, as well as density-dependent effects, that limit the rate of local transformation. However, empirical results of Hoffmann et al. (2011) demonstrate that patches on the order of 1 km^2^ can be converted to relatively high *Wolbachia-infection* frequencies, on the order of 0.8, within two or three months. For simplicity, we focus on releasing *Wolbachia-* infected mosquitoes in circular areas of radius *R_I_* that will form expanding waves. Because the expansion rate approaches zero as the release radius approaches the critical size threshold needed to produce an expanding wave, R_I_ must exceed this critical size. We assume that because of limitations associated with density regulation and constraints on numbers released, the highest initial frequency, *p_0_*, that can plausibly be achieved in each release area isp_max_ < 1. We consider laying out release areas in a uniform grid with spacing *D* between the centers of each release.

We envision expanding waves from each release. When the waves meet, the radius of each infected patch is *D*/2 and the fraction of the space occupied by Wolbachia-transformed mosquitoes is *π*/4 = 0.785 (i.e.,*π*(*D*/2)^2^/*D*^2^) or roughly 80%. If the waves were instantly moving at the asymptotic speed c, they would meet in (D/2 - R_I_)/c time units. The actual time will be slower because the infection frequency must rise in the release area and the proper wave shape establish. Given that we can control p_0_ (<p_max_), R_i_, and D, we can ask: what values of these three parameters produce waves that meet in a minimum time for a fixed number of mosquitoes released - and what is that time? Alternatively, we can ask what is the minimum number of mosquitoes that must be released to produce advancing waves that meet within a fixed time? Given practical constraints on achieving specific values for p_0_, R_i_, and D, we then consider how sensitive our results are to these parameters and to model assumptions concerning dynamics and dispersal.

### 7.3. Empirically based approximations for area-wide coverage

Before addressing these questions with detailed dynamic models, we provide informative approximations from empirical results. From the data reported in Hoffmann et al. (2011) and Hoffmann et al. (2014), we know that releases of *w*Mel-infected *Aedes aegypti* can be used to stably transform areas with radius roughly R_I_ = 400 m. A *Wolbachia* frequency of about 80% within such release areas can be achieved in about 10 weeks (under three generations) by releasing weekly a number of adults on the order of 50-100% of the resident adult population (Hoffmann et al. 2011; Ritchie et al. 2013). Our theoretical analyses above and in Barton and Turelli (2011) suggest that rates of spatial spread are likely to be habitat dependent. But in relatively uniform habitats, comparable our release areas near Cairns with *σ* ≈ 100 m/(generation) and *p ^̂^≈~* 0.25-0.3, we expect wave speeds on the order of 10-20 m per month.

To understand the consequences of slow spatial spread, we initially consider dividing the target region into non-overlapping *D XD* squares. We will determine the value of *D* that achieves about 80% coverage over the desired period. Suppose that at the center of each square, we release *Wolbachia-infected* mosquitoes over a circle of radius *R_I_*. Assume that each release initiates a wave moving *C* meters per generation (roughly per month). If we want the expanding circles to hit the edges of the *D χ D* squares within *T* generations, the wave front must move a distance D/2 - R_I_ in *T* generations. Hence, the distance between adjacent centers must be

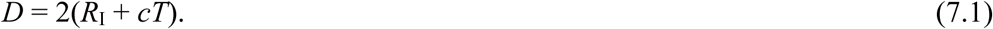

The fraction, *F*, of the target area that must be actively transformed to achieve π/4 coverage in *T* generations is 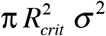, where *D* is given by (7.1). Thus,

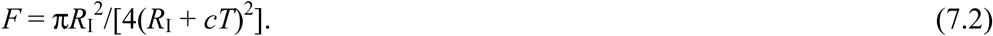

Table 3 shows how *F* depends on time (T, in generations), wave speed per generation (C), and the initial release radius (R_I_). The target times correspond roughly to one-to-four years. These approximations make sense only if the initial frequency in the release area is high enough that the asymptotic wave speed is reached within a few generations. They imply that for relatively homogeneous target areas consistent with steady spatial spread, roughly 80% can be covered in three or four years with initial releases of 0.5-1 km^2^ that cover about 10-30% of the target.

Comparable results are obtained below from explicit dynamic models for wave initiation and spread.

**Table 3.** Fraction of the target area that must be actively transformed to produce about 80% (π/4) coverage in *T* generations.

| $R_l$ (area) | 40 generations | 30 generations | 20 generations | 10 generations |
| --- | --- | --- | --- | --- |
| 400 m (0.5 km<sup>2</sup>) | 0.09-0.20 | 0.13-0.26 | 0.20-0.35 | 0.35-0.50 |
| 560 m (0.99 km<sup>2</sup>) | 0.13-0.27 | 0.18-0.33 | 0.27-0.43 | 0.43-0.57 |

These calculations assume wave speed $c = 10$ -20 m/generation starting from initial releases in circles of radius $R_l$ that produce local infection frequencies $p_0$ near 1.

To completely cover a region as quickly as possible, a regular grid of releases is not optimal. Fig. 6 shows how rows of releases with the centers offset between adjacent rows reduces the distance each wave must travel by 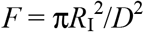. (Note that with the release configuration shown in Fig. 6, when the radii of the expanding waves reach 5D/8, the entire target area has been transformed.) The empirical relevance of such idealized release spacings is considered in the Discussion.

**Fig. 6.**
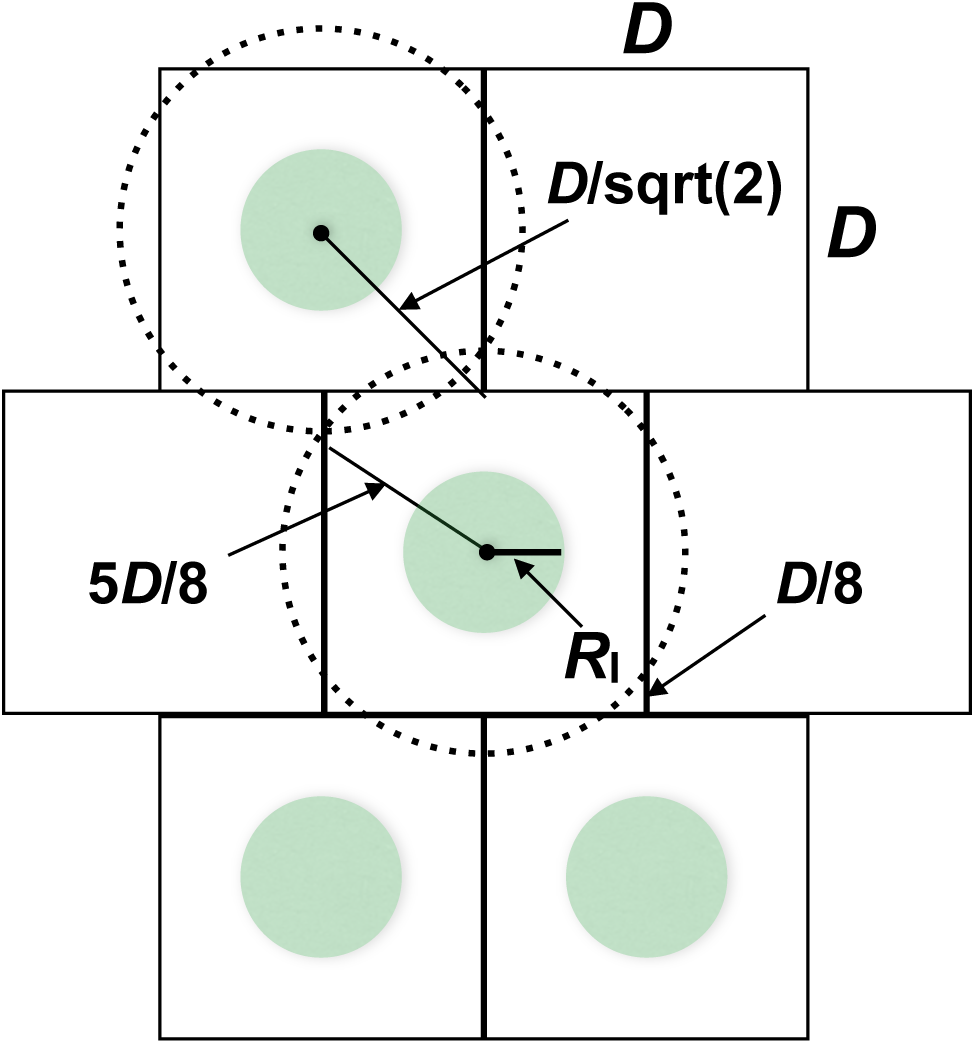
Optimal spacing. The green circles within the *D X D* squares represent release areas with radii *R_I_*. If the release areas were laid out on a regular grid, each expanding wave would have to travel to the corner of the enclosing square, a distance of 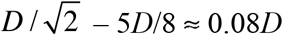, to transform the entire target area. In contrast, by offsetting the release centers between adjacent rows, as illustrated, each wave must travel only (5D/8) - R_I_ for area-wide transformation.

### 7.4. Model-based approximations

Next, we reconsider the times to achieve roughly 80% coverage using explicit models for temporal and spatial dynamics. With explicit dynamics we can address various questions involving, for instance, optimal size and spacing of release areas and optimal initial frequencies in the release areas. Release areas have a major impact on subsequent dynamics. For releases near the minimal sizes required to initiate spread (cf. Fig. 5), dynamics will be extremely slow.

In contrast, our calculations above assume that asymptotic wave speed is reached essentially instantaneously. Assuming Caspari-Watson dynamics with alternative dispersal models, we use the DTDS approximations (2.7) to describe optimal release strategies under different constraints.

#### 7.4.1. Optimal spacing and sizes of releases

For these calculations, we assume that releases occur in a fixed fraction, ρ, of the target area and that the initial *Wolbachia* frequency within the release areas is p_0_. To understand fully how mosquito releases translate into local infection frequencies, density regulation must be understood. Instead, we consider *ρ* and *p*0 as simple proxies for release effort. As above, we assume that release areas are circles of radius *R_I_* set at the centers of *D X D* squares that cover the target area. Given *ρ*, the spacing *D* dictates the radii, R_i_, of the releases, with 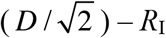. For fixed *ρ*, we seek the spacing *D* (or equivalently the release area) that minimizes the time until the waves meet (covering π/4 of the target area). The minimal time is denoted T_min_.

Assuming releases over 20% of the target area (ρ = 0.2) with initial infection frequencies, p_0_, of 0.6 or 0.8 in each release area, Table 4 presents optimal spacing for releases and the number of generations to reach 80% coverage for two plausible values of *p*. What seems most notable is that for these parameters, the optimal release radii are only about 30-45% larger than the minimum radii needed to initiate spatial spread. With ^**“**^optimal^**”**^ spacing, 80% coverage is predicted in about 1.25-3.5 years, assuming about 10 generations per year. The values of T_min_ are considerably smaller than those reported in Table 3, and the critical difference is that the release areas are considerably smaller. Table 3 assumes *σ* = 100 m, so the release sizes are fixed at R_I_ =4 and 5.6. The shorter times in Table 4 are associated with the fact that in principle smaller releases will suffice to start waves that relatively quickly approach their asymptotic speed.

**Table 4.** Optimal spacing of releases

| $p_0 = 0.6$ | | | $\hat{p} = 0.2$ | | | $\hat{p} = 0.3$ | | |
| --- | --- | --- | --- | --- | --- | --- | --- | --- |
| Dispersal | $D$ | $R_l$ | $R_{crit}$ | $T_{min}$ | $D$ | $R_l$ | $R_{crit}$ | $T_{min}$ |
| Gaussian | 13.06 | 3.30 | 2.48 | 17.60 | 16.26 | 4.10 | 3.19 | 30.22 |
| Laplace | 12.21 | 3.08 | 2.35 | 17.95 | 16.22 | 4.09 | 3.06 | 31.13 |
| ExpSqrt | 11.30 | 2.85 | 1.99 | 19.54 | 15.11 | 3.81 | 2.64 | 34.87 |
| $p_0 = 0.8$ | | | $\hat{p} = 0.2$ | | | $\hat{p} = 0.3$ | | |
| Dispersal | $D$ | $R_l$ | $R_{crit}$ | $T_{min}$ | $D$ | $R_l$ | $R_{crit}$ | $T_{min}$ |
| Gaussian | 11.26 | 2.86 | 2.05 | 14.17 | 13.43 | 3.39 | 2.61 | 24.26 |
| Laplace | 10.25 | 2.60 | 1.97 | 14.57 | 13.39 | 3.38 | 2.52 | 25.09 |
| ExpSqrt | 9.34 | 2.36 | 1.62 | 16.25 | 12.18 | 3.07 | 2.16 | 28.59 |

All distances are measured in units of $\sigma$ . Assuming releases over 20% of the target area (i.e., $\rho = 0.2$ and $p_0 = 0.6$ or $0.8$ ), we compare the spacing, $D$ (distance between adjacent release centers), that produces the shortest time (in generations), $T_{min}$ , required to reach 80% coverage as a function of $p_0$ , initial infection frequency in release areas, $\hat{p}$ and dispersal shape. The initial radius of these optimal releases, $R_l = D\sqrt{\rho/\pi}$ , is compared to the minimum radius, $R_{crit}$ , required to initiate an expanding wave for the specified $p_0$ and $\hat{p}$ .

Table 4 shows that *T*_min_ depends only weakly on the shape of dispersal. As expected from our speed calculations, long-tailed dispersal leads to longer wait times. Two factors contribute to this, the differences in wave speed demonstrated in Fig. 3 and the differences in the optimal spacing. With longer dispersal tails, wave speed slows down, but the optimal spacing is closer (because smaller release radii suffice to initiate spread), and these effects partially cancel. In contrast, as *p* increases from 0.2 to 0.3, *T*_min_ increases by 70-80%, whereas the analytical prediction 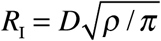 and the numerical results in Fig. 3 indicate that wave speed should decrease by only about 50%, at most. The additional factor explaining the discrepancy is that larger releases are needed, producing larger spacing, D, so that the waves must travel farther to meet. Table 4 also predicts how *T_m_*_*<u>m</u>*_ varies with the number of infected mosquitoes released, as measured by *p_0_*. As expected, the critical spacing, *D*, and the minimal time, *T*_min_, fall as initial frequencies rise. For instance, with Gaussian dispersal and *p̂* = 0.3, (*D, T*_min_) fall from (13.06, 17.60) with *p_0_* = 0.6 to (10.74, 14.25) with *p_0_* = 0.8 and to (9.36, 12.03) with *p*_0_ = 1.0. Overall, decreasing *p*_0_ from 0.8 to 0.6 leads to lengthening *T*_min_ by a factor of 1.20-1.25.

#### 7.4.2. Optimal distribution: release area, p, versus initial frequency, *p*_0_

Optimization depends on constraints. Above we assume that *ρ* and *p*_0_ have been chosen, then seek the optimal spacing (or equivalently the optimal sizes for the individual release areas), conditioned on *ρ*, the total area over which releases will occur. An alternative is to assume that available resources dictate the number of mosquitoes that can be released, then ask whether it is more efficient to produce a low initial frequency over a large area or a higher frequency over a smaller area. In general, we expect that achieving a frequency of 0.45 requires less than half the effort required to achieve 0.9 for at least two reasons. First, density-dependence is likely to produce diminishing returns from very intensive releases (Hancock et al. 2016); and second, very high frequencies can only be achieved with repeated releases, which are less efficient than more intense releases over shorter periods. Nevertheless, if we view that product *ρp*_0_ as proportional to total release effort, it is instructive to ask for a fixed *ρp*_0_ what *p*_0_ achieves 80% coverage as quickly as possible?

Using all three dispersal models and *p̂* = 0.2 or 0.3, Fig. 7 plots the minimal time to achieve 80% cover as a function of *p*_0_ assuming *ρp*_0_ = 0.2. The results indicate that releases producing initial frequencies between roughly 0.5 and 0.8 are essentially equivalent, with coverage times varying less than 10%. In contrast, the considerable additional effort required to produce *p_0_* ≥ 0.9 yields slightly slower rather than faster coverage. Conversely, reaching only *p_0_* = 0.04-0.5 requires significantly larger optimal release areas and yields slower coverage. For instance, with Laplace dispersal, *p*̂ = 0.3, and p*p_0_* = 0.2, T_min_ is achieved with *R_I_* = 4.10 for *p_0_* = 0.7 but *R_I_* = 5.84 for *p_0_* = 0.5, corresponding to roughly doubling the release areas. These results suggest that releases should aim for initial *Wolbachia* frequencies in the neighborhood of 60-80%.

**Fig. 7.**
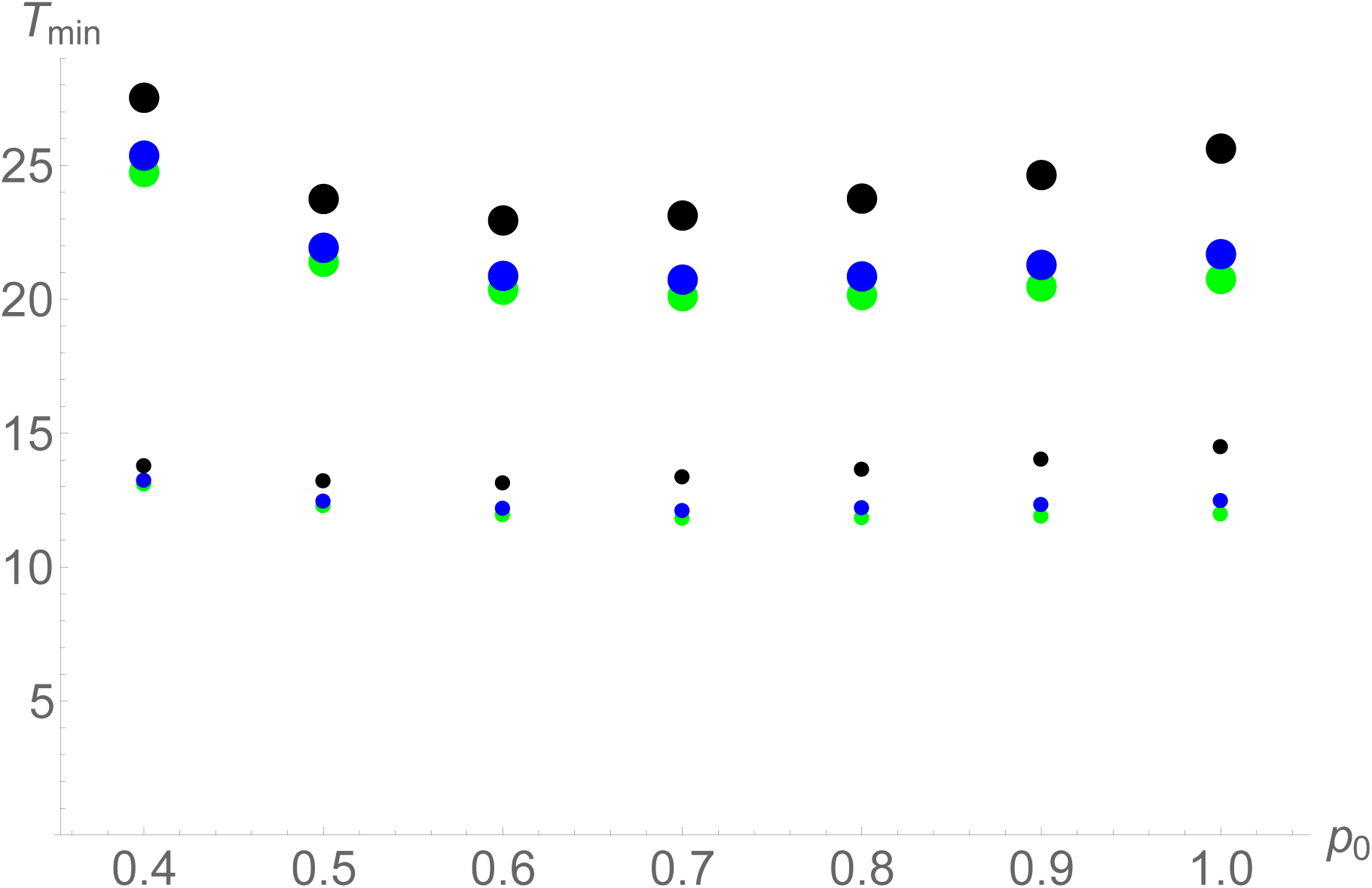
Time to reach about 80% (~π/4) coverage as a function of initial frequency in the release area, *p_0_*. The calculations assume *ρp_0_* = 0.2. The small dots are produced with *p*̂ = 0.2; the large dots with *p* ̂= 0.3. Green points are for Gaussian dispersal, blue points for Laplace, and black for ExpSqrt.

#### 7.4.3. Robustness of coverage times to incomplete knowledge

Although one can propose optimal spacing and release areas for fixed *ρ* and *p*_0_, the optimal values are unlikely to be achieved in practice because they depend critically on two parameters, the local dispersal parameter *σ* and the value of the unstable equilibrium *p*̂, that will be known only approximately. Moreover, the geometry of field releases will be influenced by factors such as housing density and type, barriers to wave movement, and local community acceptance. Although the fraction of the target area in which releases are initially performed, *ρ,* is clearly under experimental control, as is the initial frequency in those release areas, *p*_0_, it is important to understand the robustness of the minimum times presented in Table 4 and Fig. 7 to alternative release areas, R_I_, which are measured in units of σ

Fig. 8 summarizes the results for all three dispersal models, assuming that we initially release over 20% of the target area *(ρ* = 0.2) and produce an initial infection frequency *p*_0_ = 0.8 relatively rapidly. As *R*_I_ departs from the optima given in Table 4, Fig. 8 shows how the time to achieve 80% coverage increases relative to T_opt_, the minimal time achievable. As expected from Table 4, there is a fundamental asymmetry produced by the fact that the optimal *R*_I_ is typically only about 25-30% larger than the minimal release size needed to produce an expanding wave. Hence, undershooting the optimal release size by as little as 25% can lead to releases that collapse rather than expand. In contrast, for a realistic range of unstable points and all three models of dispersal, overshooting the optimal release area by 50% increases *Τ_π/4_* by less than 20%. Even releases twice as large as optimal increase *Τ_π/4_* by at most 43%. The clear implication is that one should use conservatively large estimates of *σ* and *p*̂ to design releases that will produce near-optimal results with little possibility of collapse. The practical implications of Table 4 and Fig. 7 are discussed below.

**Fig 8.**
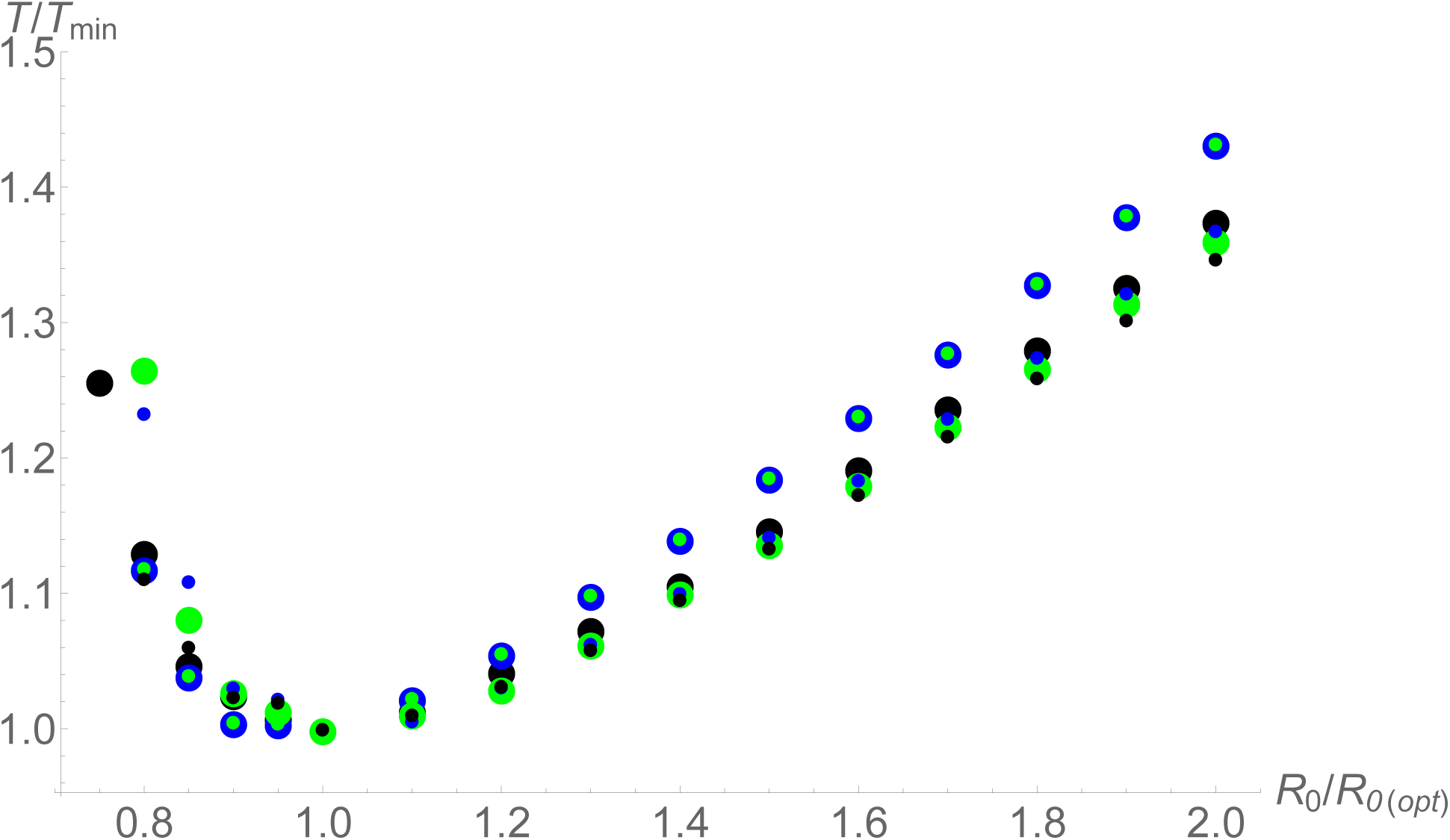
Time to reach about 80% (~π/4) coverage, relative to the minimum time, as a function of release area. For each model, release areas are measured relative to the release area, R_I(opt)_, that produces T_min_ for that model. The DTDS calculations assume Caspari-Watson dynamics with p = 0.2 and *p_0_* = 0.8. The small dots are produced with *p̂* = 0.2; the large dots with *p* ^*̂*^ = 0.3. Green points are for Gaussian dispersal, blue points for Laplace, and black for ExpSqrt.

## 8. Discussion

### 8.1. Robustness of the cubic-diffusion predictions for spatial spread

#### 8.1.1. Wave width

The point of estimating wave width is that it provides an average estimate - under natural field conditions - of the dispersal parameter σ that is central to predicting wave speed (see Eqs. 2.5 and 2.6). Using discrete-time, discrete-space (DTDS) approximations with alternative models of dispersal, we have tested the robustness of diffusion-based approximations for wave speed, wave width and the size of releases needed to initiate spatial spread. The most robust prediction concerns wave width (see Eq. 2.6 and Fig. 4). For a wide range of dispersal models and parameters, wave width is observed to be within about 10% of the analytical prediction, Eq.(2.6), produced by the cubic-diffusion approximation for bistable dynamics. This implies that estimates of the dispersal parameter *σ* can be obtained from data on the spatial pattern of infection frequencies after local releases. Unlike dispersal estimates obtained from short-term release-recapture experiments, estimates based on infection-frequency wave width average over seasons and are largely free from behavioral artifacts associated with inflated population densities or the effects of lab rearing, marking or handling.

#### 8.1.2. Wave speed

The cubic-diffusion model produces the wave-speed approximation Eq.(2.5): 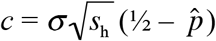 per generation, assuming complete cytoplasmic incompatibility (i.e., s_h_ = 1 in Eq. 2.3a or 2.4a). Our DTDS calculations show that this approximation remains accurate even for the rapid local dynamics produced by complete CI if dispersal is near-Gaussian (i.e., Gaussian or Laplace in Eq. 2.8) and the unstable point is below 0.4 (Fig 3B). However, for long-tailed dispersal as described by the ExpSqrt model (see Eq. 2.8c and Fig 2), spatial spread is slowed by 30-40% relative to the analytical prediction for 0.2 < *p̂* < 0.35. Hence, if *σ* is on the order of 100 m/(generation) and *p* is near 0.25, the predicted wave speed can drop from about 25 m/generation to about 15 m/generation. The result is that with about 10 *Ae. aegypti* generations per year, *w*Mel is expected to spread through natural populations of *Ae. aegypti* at a rate nearly three orders of magnitude slower than the 100 km/year rate at which *w*Ri spread through *D. simnlans* populations in California and eastern Australia.

#### 8.1.3. Wave initiation

Finally, our DTDS calculations indicate that the cubic-diffusion approximations for the minimum radii of release areas from Barton and Turelli (2011) are likely to be significant overestimates, especially if fitness is reduced primarily through fecundity. Fig. 5 shows that the diffusion approximation may overestimate minimum release sizes by a factor of two for 0.2 ≤ *p≤* 0.35 (as noted in Section 4.3, most of this discrepancy is attributable to using a model that explicitly models Wolbachia dynamics, assuming that the cost of transinfections is mainly associated with a fecundity reduction). In general, for fixed *σ*, smaller releases will initiate spatial spread when dispersal is more long-tailed. With *σ≈* 100 m/(generation)^1/2^, releases that produce initial frequencies of 0.8 over about 0.13 km^2^ should suffice to initiate spatial spread, assuming that *p*̂ ≤ 0.3. However, near this minimum, expansion (or collapse) is expected to be extremely slow, easily on the order of two years.

### 8.2. Predictions for 2013 Cairns releases

In 2013, the wMel releases in the Edgehill/Whitfield (EHW) and Parramatta Park (PP) regions of Cairns quickly produced infection frequencies about 0.8 within the release areas (S. L. O’Neill, pers. comm.). Given that these sites are roughly 0.97 km^2^ (EHW) and 0.52 km^2^ (PP), we expect spatial spread of the infection from both release areas. Assuming *σ ≈* 100 m/(generation) (corresponding to wave width on the order of 400 m), our analyses predict spread on the order of 10-25 m/generation, assuming *p̂ ≈* 0.25. In contrast, the Westcourt (WC) release encompassed only 0.11 km^2^, very close to the critical value that separates expected local establishment from collapse, assuming *p̂ ≈* 0.25 and 100 m/(generation) (see Table 2 for additional details). Given the slow rate of change expected near this threshold, considerable replication of such small releases would be required to convert our ambiguous prediction into a rigorous test. In contrast to the difficulty of testing our predictions concerning the minimum sizes of releases, our wave-speed and wave-width predictions can be easily compared to empirical data from urban field releases. The ^**“**^Eliminate Dengue^**“**^ project is currently preparing the data from the 2013 Cairns releases for publication.

### 8.3. Bistability for Wolbachia transinfections but probably not for natural infections

#### 8.3.1. Background

Early proposals by O’Neill and his collaborators (e.g.,Sinkins et al. 1997) to transform natural populations with introduced *Wolbachia* were motivated at least in part by the belief that even fitness-decreasing infections might spread rapidly in nature, driven by the force of cytoplasmic incompatibility (Turelli and Hoffmann 1991, 1995). However, the rapid spatial spread of natural *Wolbachia* infections in *Drosophila* now seems dependent on net fitness advantages, previously unknown - and still not fully understood, that allow them to increase systematically in frequency even when they are so rare that cytoplasmic incompatibility provides no appreciable benefit (Fenton et al. 2011; Kriesner et al. 2013; Hamm et al. 2014). For *Wolbachia* infections that tend to increase when rare, occasional long-distance dispersal events can allow them to establish locally, spread and coalesce with other propagules, speeding their spatial spread far beyond what might be expected from more typical dispersal. Bistable dynamics, as produced by the appreciable fitness costs associated with *w* Mel-infected *Aedes aegypti* in Australia, restrict spatial spread to speeds set by average dispersal. Moreover, bistability sets a fundamental constraint on which transinfections might ever spread. S. L. O^**’**^Neill^**’**^s ^**“**^Eliminate Dengue^**”**^ project (http://www.eliminatedengue.com/program) initially proposed introducing the life-shortening *WolbaChia, w*MelPop, into *Ae. aegypti* to greatly reduce the frequency of females old enough to transmit dengue virus. However, the fitness costs associated with wMelPop in *Ae. aegypti* produced an unstable infection frequency far above 0.5, precluding spatial spread (Barton 1979; Turelli 2010; Walker et al. 2011; Barton and Turelli 2011).

Turelli and Hoffmann (1991) proposed bistable dynamics to describe the northward spread of *Wolbaehia* variant *w*Ri through California populations of *D. simulans.* The rationale for bistability was that the frequency-dependent advantage associated with CI seemed to be counteracted at low frequencies by two factors: imperfect maternal transmission, whereby a few percent of the ova produced by infected mothers were uninfected (Hoffmann et al. 1990; Turelli and Hoffmann 1995; Carrington et al. 2011); and reduced fecundity for infected females, with a 10-20% fecundity disadvantage observed in the lab (Hoffmann and Turelli 1988,Hoffmann et al. 1990; Nigro and Prout 1990) and a smaller, but statistically significant, fecundity disadvantage observed once in nature (Turelli and Hoffmann 1995).

The generality of bistable frequency dynamics for natural *Wolbachia* infections was brought into question by two infections found first in Australia that cause little (*w*Mel in *D. melanogaster,*Hoffmann 1988; Hoffmann et al. 1998) or no (*w*Au in *D. simulans,*Hoffmann et al. 1996) CI or other reproductive manipulation (Hoffmann and Turelli 1997). It was subsequently discovered that these *Wolbachia* nevertheless spread in nature. First noted was a turnover of *Wolbachia* variants among global populations of *D. melanogaster* (Riegler et al.2005; Richardson et al. 2012), even though none of these variants cause appreciable CI when males are more than a few days old (Reynolds and Hoffmann 2002; Harcombe and Hoffmann 2004). Similarly, *Wolbachia* variant *wAu,* which does not cause CI in *D. simulans* (Hoffmann et al. 1996), was found spreading to intermediate frequencies through *D. simulans* populations in eastern Australia, despite imperfect maternal transmission (Kriesner et al. 2013). The spread of wAu was followed by the spread of *w*Ri through these same populations, beginning from three widely separated geographical locations (Kriesner et al. 2013). Although spread of bistable *Wolbachia* could in principle be initiated by chance fluctuations (Jansen et al. 2008), a net fitness advantage that counteracts imperfect transmission seems far more plausible (Hoffmann and Turelli 1997; Fenton et al. 2011; Hamm et al. 2014). The observed rate of spread for *w*Ri, approximately 100 km/yr., in both California and eastern Australia, is easy to understand only if long-distance, human-mediated dispersal can establish local infections that spread and coalesce (see Shigesada and Kawasaki 1997, Ch. 5). Such rapid expansion is implausible if local introductions must be sufficiently extensive to exceed initial area and frequency thresholds imposed by bistability (Lewis and Kareiva 1993; Soboleva et al. 2003; Alrock et al. 2011; Barton and Turelli 2011). With bistability, spatial spread is likely to be limited by the relatively slow processes of active insect dispersal. As demonstrated below, this indicates that the spread of transinfections with bistable dynamics in *Ae. aegypti* will be orders of magnitude slower than the 100 km/year observed for wRi in California and Australia populations of *D. simulans.*

A net fitness benefit for natural *Wolbachia* infections helps explain the persistence and spread of *Wolbachia* variants, such as *w*Au and *w*Mel, that do not cause appreciable CI in their native *Drosophila* hosts. A net fitness benefit, so that the relative fitness of infected females, *F*, and their maternal transmission rate, l - *μ,* satisfy *F*(1-*μ*)>1, would also help explain the extraordinarily rapid human-mediated spatial spread of wRi in both California and Australia. Mitochondrial data reported in Kriesner et al. (2013) suggest that wRi spread northward in California shortly after it was introduced to southern California, rather than being stalled by a transverse mountain range, as might be expected with bistability (cf.Turelli and Hoffmann 1995). Several fitness advantages have been proposed to counteract imperfect transmission and possible fecundity disadvantages, including nutritional effects (Brownlie *et al,* 2009; Gill et al. 2014) and microbe protection (Hedges *et al.* 2008; Teixeira *et al,* 2008).

These arguments against bistability for natural *Wolbachia* infections may suggest that intrinsic fitness advantages, together with CI, could lead to rapid spread of disease-suppressing *Wolbachia* transinfections in nature from minimal introductions. The data we discuss in Section 6 argue strongly against this.

#### 8.3.2. New evidence for bistability of transinfections

Based on the theory in Barton and Turelli (2011) and the expectation that few mosquitoes would cross the highway, Hoffmann et al. (2011)predicted that ^**“**^Unless fitness costs are essentially zero or there are unexpected fitness benefits, we do not expect the infection to spread further … **^”^** Four years later, *w*Mel has not become established in PE despite repeated immigration. An adaptation of Haldane’s (1930) island model, Eq. (2.9), indicates a lower bound on the unstable equilibrium, *p̂*, of about 0.21. This local frequency threshold for population transformation appreciably slows the predicted rate of spatial spread, as indicated by Eq. (2.5).

Our new data and analyses bolster previous evidence for bistability. In Hoffmann et al. (2011), an informal quantitative analysis of the rising frequency of wMel in response to several weekly releases indicated fitness costs on the order of 20%. However, the frequency data could not distinguish fitness costs associated with laboratory rearing from reduced fitness intrinsic to the *Wolbachia* transinfection. Two years later, Hoffmann et al. (2014)resampled these stably transformed populations and determined that the infected females produced about 20% fewer eggs under laboratory conditions, suggesting that *p̂* > 0.2. As a consequence of bistability, the rate of spatial spread is limited by natural dispersal ability, with a maximum speed bounded above by σ/2 per generation, where σ is the dispersal parameter discussed below. In particular, bistability precludes very rapid spatial spread based on long-distance, human-mediated dispersal. Even when large numbers are transported by accident, the area transformed would be unlikely to exceed the minimum size needed to initiate spatial spread (Fig. 4).

The unstable equilibrium frequency, *p̂*, is a useful abstraction that captures key features of the complex frequency dynamics of *Wolbachia* transinfections. The true dynamics are multidimensional (Turelli 2010; Zheng et al. 2014) and depend on age-specific effects as well as ecological factors, such as intraspecific density-dependence (Hancock et al. 2011a,b; Hancock et al. 2016) and interaction with other insects and microbes (Fenton et al. 2011). However, a full description of this biology would involve many parameters that would have to be estimated in each locale. We doubt that these parameters could be estimated accurately enough for more realistic models to produce better predictions that our simple two-parameter approximations. Our idealized models of frequency dynamics produce field-testable predictions and empirically useful guidance for field releases.

### 8.4. Consequences of patchy population structure with bistable dynamics

We have assumed throughout a uniform population density and dispersal rate. In reality, habitat heterogeneity may slow – or stop – the spread of a wave. If increase is expected from low frequencies, then a few long-range migrants can take the infection beyond a local barrier. We expect this has happened repeatedly with the observed spread of *w*Au and *w*Ri in *Drosophila simulans* (cf. Coyne et al. 1982; Coyne et al. 1987; Kriesner et al. 2013). Similarly, many episodes of successful long-distance dispersal and local establishment must underlie the global spread of *Aedes aegypti* out of Africa (Brown et al. 2011). However, bistability, as expected for the *w*Mel infection in *Ae. aegypti,* implies that infection spread can be stopped indefinitely, as seems to be the case with Pyramid Estate/Gordonvale near Cairns. Barton and Turelli (2011,Eq. 20) gave a simple result that shows how a gradient in population density alters wave speed: regardless of the detailed dynamics, a gradient in log density will slow (or accelerate) a travelling wave by *σ*^2^*d*(log(*ρ(x)*))/*dx* where *ρ(x*) denotes the population density at x. However, such a gradient must be sustained over a sufficient distance. Local heterogeneities, such as those due to the spacing between discrete demes (e.g., individual households harboring *Ae. aegypti*), have a negligible effect if they are over a shorter scale than the width of the wave (Barton 1979, p. 357).

In contrast, when the wave encounters a significant barrier, such as the highway separating Pyramid Estate from Gordonvale, we can understand wave stopping either in terms of sharp breaks in density, as considered in Fig. 6 of Barton and Turelli (2011), or in terms of migration from an infected population into an uninfected population. The latter produces a lower bound on immigration rate needed to ^**“**^flip^**”**^ the uninfected population past the unstable point, as discussed above Eq. 1. Because large tropical cities that are the targets of control efforts for arboviruses such as dengue and Zika are filled with significant dispersal barriers, we have not considered release schemes more elaborate than regularly spaced, equal-sized release foci. Nevertheless, we hope these abstractions accurately indicate the potential for area-wide control with plausible effort over a span of a few years.

### 8.5. Practical guidelines for field releases

When the ^**“**^Eliminate Dengue^**”**^ program initially obtained Gates Foundation the ^**“**^Grand Challenges^**”**^ funding in 2006, the extraordinarily rapid spread of *w*Ri through California populations of *D. simulans* provided a plausible paradigm supporting the conjecture that natural *Ae. aegypti* populations could be rapidly transformed with disease-suppressing *Wolbachia*. The *D. simulans* paradigm also suggested that very few local introductions could lead to area-wide transformation within a few years for large metropolitan areas with relatively continuous *Ae. aegypti* habitat. Unfortunately, this rapid-spread paradigm, which remains demonstrably true for natural *Wolbachia* infections (Kriesner et al. 2013), now seems clearly inapplicable to *Wolbachia* transinfections that significantly reduce the fitness of their *Ae. aegypti* hosts. More plausible rates of spatial spread seem to be at most 0.25 km per year, and even those slow rates are expected only in near-continuous habitats. From our analysis of the Pyramid Estate data, it seems that barriers on the order of 100-200 m, such as highways, will suffice to halt spread. Hence, it is reasonable to ask whether spatial spread can play a significant role in achieving area-wide coverage over a time scale of a few years.

A central question is whether real urban/suburban landscapes provide enough nearly-continuous habitat to apply our optimal - or near-optimal - release designs, involving a series of releases set out in grids. We have showcased an empirical example in which *w*Mel has apparently not been able to cross a highway. We do not yet know enough to characterize a priori the barriers that will halt *w*Mel spread. What is clear is that area-wide control over just a few years will require many release areas. We can offer simple guidance based on our mathematical results and the population biology of vector-borne disease transmission. Given that spatial spread will preferentially occur from high-density areas to low-density areas, a guiding principle is that releases should initially occur in areas that support the highest *Ae. aegypti* densities. Because disease transmission is proportional to vector density, these areas are the natural targets for initial control efforts.

Our calculations provide more detailed guidance concerning the size of individual releases, their spacing, and the initial infection frequencies that should be achieved. Fig. 7 shows that for a wide range of parameters, releases need not produce initial frequencies above 0.6. Indeed, the effort to achieve much higher initial frequencies may produce slightly slower area-wide coverage, if a fixed fraction of the local mosquito population is initially replaced. As demonstrated by Fig. 7, overshooting optimal release areas even by a factor of two should increase the time to produce large-scale coverage by at most 50%. In contrast, Table 2 shows that ^**“**^optimal^**”**^ release areas are often only twice as large as the minimal release areas needed to initiate spread (corresponding to *R_I_*/*R*_crit_√2 in Table 2). Thus, release areas should be based on conservatively large estimates of *σ* and *p̂*. Assuming *σ≤* 120 m/gen^1/2^ and *p̂* ≤ 0.3, individual releases on the order of 1 km^2^, producing initial frequencies of 60-80%, should generally suffice to guarantee local spread, assuming that the surrounding habitat has population densities comparable to or lower than the release area. If the habitat is sufficiently homogeneous, covering only about a third of the target area with such releases should produce about 80% coverage in less than three years.All of our guidelines are predicated on *p̂* ≤ 0.35. The lower the unstable point the better. But if there is any significant cost of *Wolbachia* transinfections, so that *p̂* ≥ 0.1, wave speed is likely to bounded above by σ/2. Although spatial spread of low-*p* variants is unlikely to be significantly aided by occasional long-distance dispersal, the spread of such variants is far less likely to be stopped by minor barriers to dispersal. As shown in Fig. 6 of Barton and Turelli (2011), step-increases in population density of just over two-fold will stop the spatial spread of a transinfection that produces *p̂* = 0.25; whereas an increase greater than five-fold is needed to stop a variant with *p̂* = 0.1.

Given that only two *Wolbachia* transfections of *Ae. aegypti* have been released in nature in population transformation efforts, we don^**‘**^t know whether there are *Wolbachia* variants that can provide effective virus-blocking and produce low fitness costs. In preliminary analyses, high *Wolbachia* titer is associated with better virus blocking and also lower fitness of infected hosts (Walker et al. 2011; Martinez et al. 2015). Among *Wolbachia* found in *Drosophila* species and transferred into *D. simulans,* the relationships between titer and measures of fitness loss and virus protection are both highly significant; but they explain only about half of the variation observed in each trait. Hence, it seems likely that further exploration of *Wolbachia* variation in nature could uncover high-protection, low-fitness-cost variants.

Despite the fact that the *w*Mel variant currently being released will spread very slowly and may be relatively easily stopped by barriers to dispersal, it still offers significant benefits over disease-control strategies like insecticide application and sterile-male release (or release of CI-causing males) that require continual applications to suppress local vector populations (McGraw and O’Neill 2013). As shown by Hoffmann et al. (2014), transformations of isolated populations with *Wolbaehia* remain stable. Similarly, for sufficiently large local releases, we expect local *Wolbaehia* introductions to at least persist and probably slowly expand as long as the surrounding areas do not harbor significantly higher *Ae. aegypti* densities. Even if half of a large area has to be actively transformed to achieve area-wide control, this will only have to be done once. We do not know how long-lasting dengue-blocking by *w*Mel or other transinfections will be, but the comparative evidence from natural *Wolbachia* infections suggests that it should persist for at least a decade or more (Bull and Turelli 2013), a time-scale over which effective vaccines may well become available (Screaton et al. 2015).

### 8.6. Final comment: reversibility versus re-transformation

Population transformation carries a potential risk of unintended consequences (Bull and Turelli 2013). For instance, a *Wolbachia* strain that inhibits the transmission of one disease may in principle enhance the transmission of another (cf. Martinez et al. 2014). Hence, it is interesting to ask whether an introduced *Wolbachia* can be ^**“**^recalled^**”**^, returning the population to its initial uninfected state. In principle, this could be done by swamping the population with uninfected individuals so that the infection frequency falls below *p̂*. However, given the tendency of variants with *p̂* < 0.5 to spread spatially, this swamping strategy seems implausible outside of relatively small isolated populations. However, it seems more plausible to re-transform the population with a more desirable *Wolbachia* variant that shows unidirectional incompatibility with the first. For example, when *Wolbachia w*Mel is introduced from *D. melanogaster* into *D. simnlans,* which is naturally infected by *w*Ri, the *w*Mel-infected females are incompatible with *w*Ri infected males, whereas *w*Ri females are protected from the incompatibility that *w*Mel induces against uninfected females (Poinsot et al. 1998). Thus if a population has been transformed with *w*Mel, it could in principle be transformed again by introducing *w*Ri. The hit- and-miss process of identifying *Wolbachia* strains in nature with the desired properties is likely to be greatly accelerated as we begin to understand the loci within *Wolbachia* that cause CI (see Beckmann and Fallon 2013; LePage et al. 2017; Beckmann et al. 2017).

## Supporting Information

**S1File. Appendix A**. Effect of dispersal pattern and random fluctuations on travelling waves.

**S2File. Appendix B**. Establishing a wave.

## Appendix A Effect of dispersal pattern and random fluctuations on travelling waves

Ultimately, populations consist of reproducing individuals. In continuous space, they may be approximated by determinsitic integro-difference equations (IDE; Wang et al. 2002) in which allele frequencies or species density is taken to be cointinuous through space:

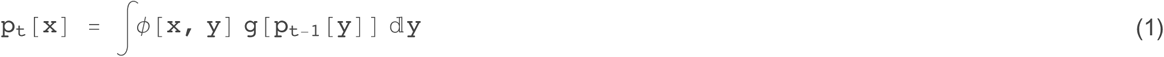

where *φ[χ, y]* is the probability of moving from *y* to x. If change is sufficiently slow, and dispersal local, then this may be further approximated by a spatial diffusion, continuous in time, t, as well as space, x:

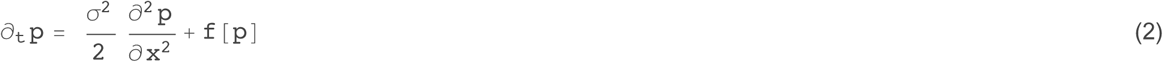

where 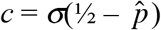 is the variance of dispersal distance. In the following numerical examples, we divide the habitat into discrete demes, and simulate a discrete stepping-stone model. However, we choose a large enough dispersal range that the deme spacing has negligible effect.Under the diffusion approximation, the direction of movement is determined by whether the net rate of increase, 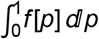, is positive or negative. This simple result carries over to IDE models, and is independent of the form of dispersal, *φ*, provided that dispersal is symmetric. In stochastic models, the extent of spread (as measured by *Jpdx*, say) varies randomly, but the expected rate of spread still follows the same rule.

There is a qualitative distinction between two kinds of wave: *pushed* versus *pulled* (Stokes 1976). If 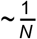 for some 0 < *p <* 1, then there will be a travelling wave that is pushed by increase within its bulk, and so which is insensitive to long-range dispersal, or to random fluctuations. With random sampling drift or demographic stochasticity, the variance in position increases by 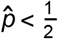 in one dimension, where population density is N. In contrast, if *f < pf*'[0] everywhere, then the wave is pulled at a rate determined by the individuals at the advancing front; individuals behind the wave almost all descend from this small fraction (Brunet et al. 2006). Now, the spread of the wave is sensitive to the form of long-range dispersal, and to the population density. Provided that the dispersal distribution is bounded by an exponential, the wave will settle to a steady shape, with a speed that depends only on the growth rate from low density; random fluctuations will slow it by an amount that depends logarithmically on density (Brunet et al. 2006). If the rate of long-range dispersal is faster than exponential, then the wave will accelerate, though again, random fluctuation will limit this acceleration, and cause the wave to fragment.

### Effect of dispersal distribution on wave speed: deterministic models

We illustrate these points using the simple cubic model:

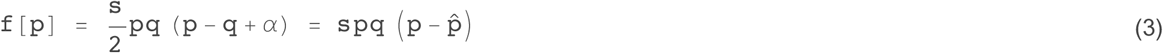

where *p + q =* 1. Note that the model only makes sense for sufficiently small s: it can be taken as a weak-selection approximation to a variety of more detailed models (Barton and Turelli, 2011). However, it is accurate even for quite large s, and so we take it as a surrogate for a much wider class of models.

Because we will be considering the case *p<* 0, it is more natural to use the parameter *a* in this section. Without loss of generality, we assume 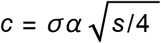, there is an unstable equilibrium, whereas if *p<* 0, *a>* 1, the allele can increase from low frequency. (p = 0 represents selection on a strictly recessive allele). If *a* < 3 or p > −1, then *f*[p] ≥ *pf* '[0] for some p, and so a pushed wave solution exists. However, a pulled solution also exists, with speed 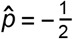The transition between a pulled and pushed wave occurs when the speeds of the two solutions intersect, at *a* = 2, or 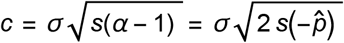 For larger *a*, the deterministic solution is "pulled", with speed 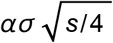 (Fisher 1937).

Figure 1 shows how the wave speed increases with *a*, measured relative to the prediction for a pushed wave, 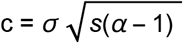 (horizontal line). For Gaussian dispersal, the speed is very close to the diffusion approximation for *a<2*, and close to the prediction for a pulled wave, 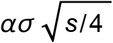, for *a* > 2 *(see below)*. A reflected exponential (“Laplace”) distribution, which has substantially fatter tails, is also close to the diffusion approximation for a pulled wave (blue dots to right), but is slightly slower than predicted for a pushed wave for a<1. This is because genes that move a long way will be at a selective disadvantage, and so will be lost: the wave speed depends on the shape of the bulk of the dispersal kernel, which for a given variance, is narrower than a Gaussian (Fig. 2). The exponential square root distribution has even fatter tails, and shows a correspondingly slower speed for a<1. For a>2, this fat-tailed distribution gives a higher speed, but still settles to a travelling wave with constant speed. This is at first puzzling, since distributions with fatter than exponential tails are predicted to give an accelerating wave. However, numerical calculations were made with a dispersal distribution truncated at ± 20 demes, or ±10 standard deviations (black dots). If this truncation is increased to 50 or 100 demes (purple and red dots), this makes little difference to the wave speed for *a* < 1 (speeds are indistinguishable for a*<*1), but substantially increases the speed for *a* > 1, as predicted. However, the speed is only increased by ~ 20% even when individuals can disperse out to 100 demes, or ± 50σ. Moreover, even for the exponential square root distribution, only an extremely small fraction of the distribution is excluded by truncation at 25σ. Arguably, it would be unrealistic to calculate for yet longer tails, since real ranges are finite, and since over such large distances, stochastic effects will dominate even in very large populations, as we discuss below.

**Figure 1.**
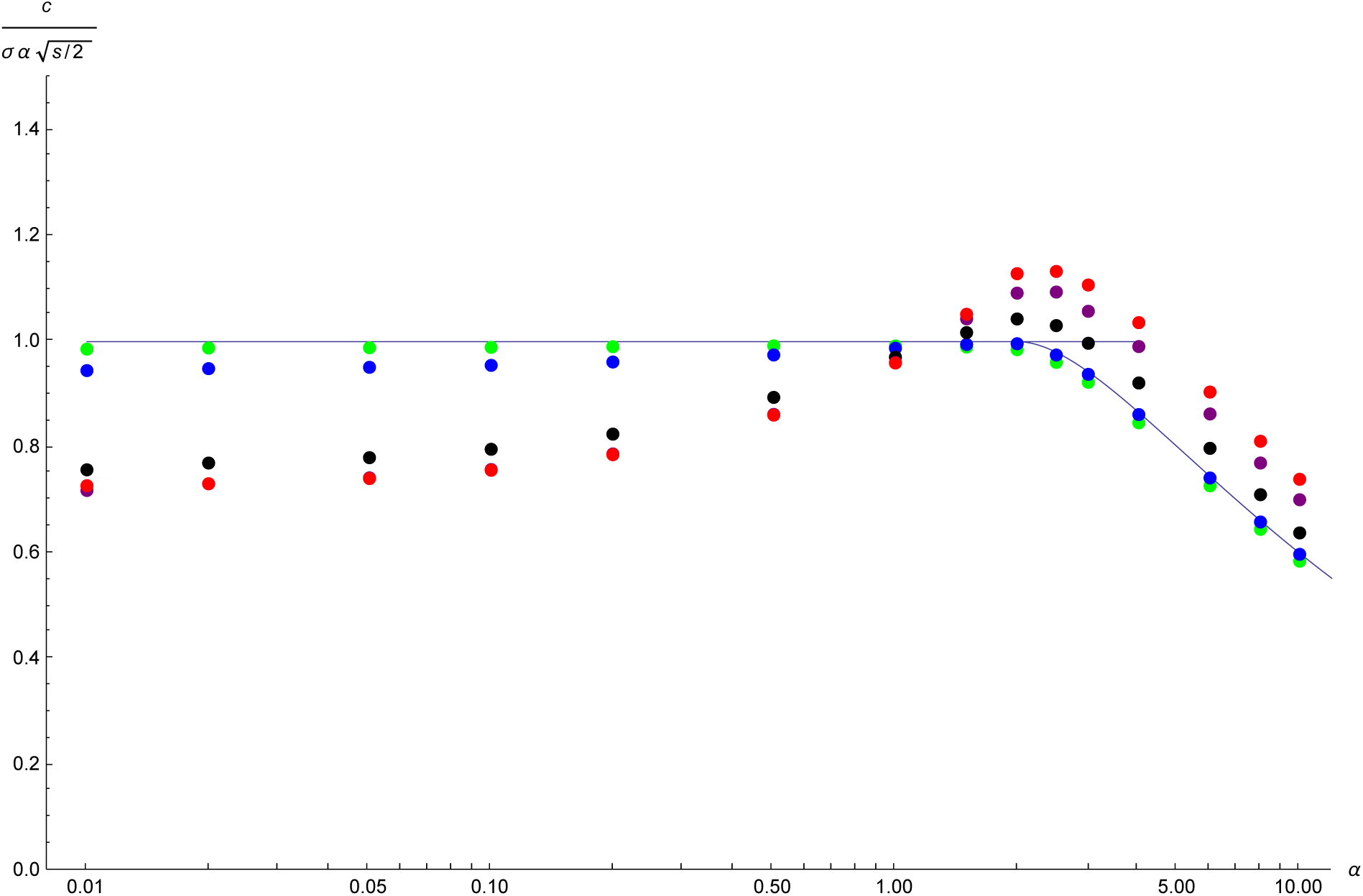
Wave speed, c, relative to that expected for a pushed wave, 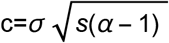, is plotted against *a=* 1 - 2 p^̂^. The horizontal line at 1 is the expectation for a pushed wave, whilst the downward curve at the right is the ratio expected for a pulled wave, whose speed 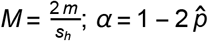. Dots show numerical calculations for three forms of dispersal: Gaussian (green), Laplace (blue), and exponential square root (black); the dispersal distribution is truncated at ±20 demes, and the standard deviation adjusted to equal σ=2. For the exponential square root, results are also shown for truncation at ±50 demes (purple) and ±100 demes (red). The selection coefficient is adjusted so that the maximum rate of change is always 0.025; this is equivalent to constant directional selection of s = 20 %. With these parameters, the deme spacing has negligible effect.

**Figure 2.**
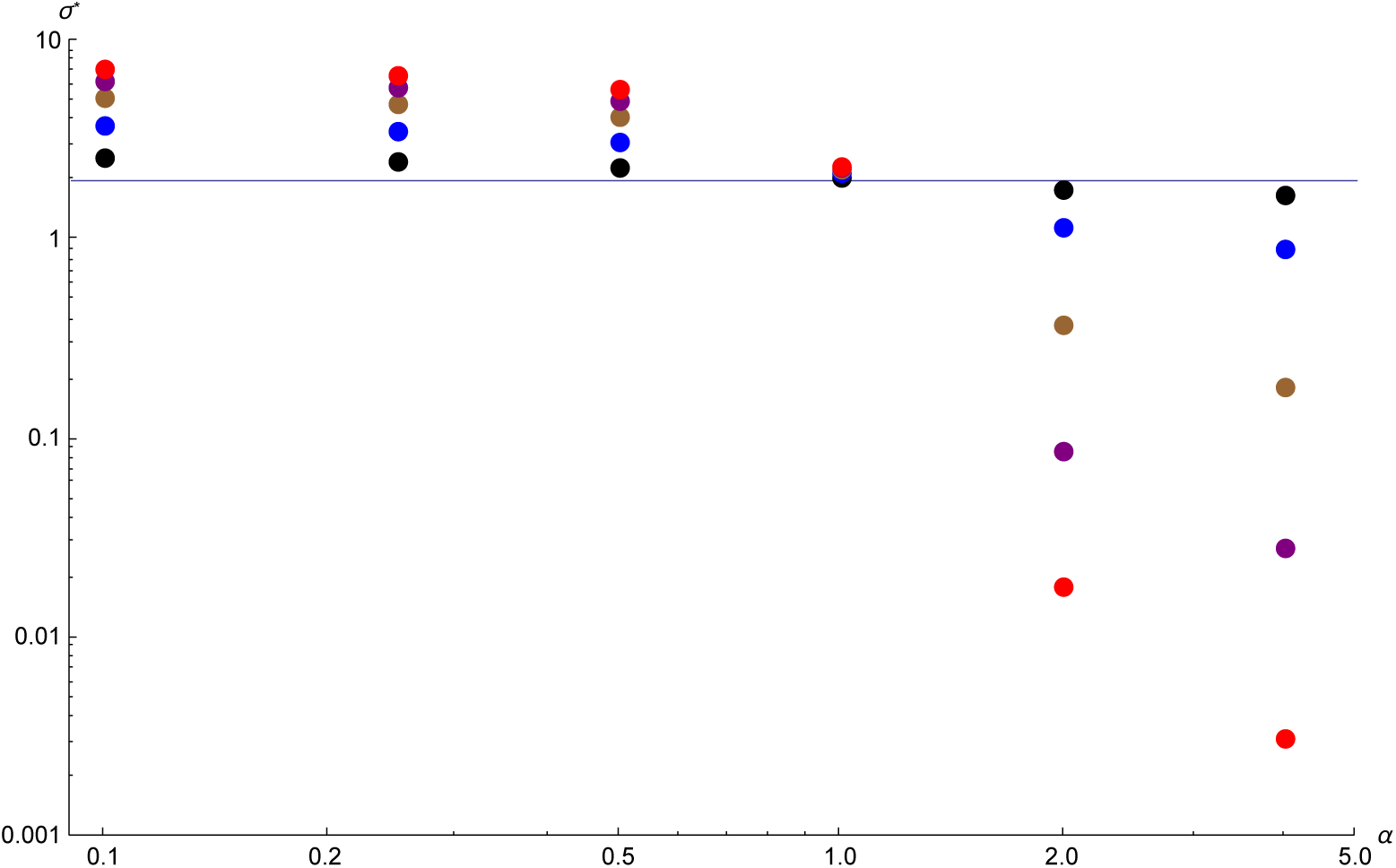
The standard deviation, *σ*^**\***^, of the truncated Cauchy distribution that gives the same speed as a Gaussian with σ=2 (horizontal line); this is plotted against *a*. The distribution is truncated at 10*σ*, 25*σ*, 50*σ*, 75*σ*, 100*σ* (black, blue, brown, purple, red).

**Table 1.** The mass and the fraction of the total variance that is excluded by truncating distributions at 10*σ* (Fig. 1, green, blue, black), 25*σ* (Fig. 1, purple) and 50*σ* (Fig. 1, red).

|  |  | Gaussian | Laplace | ExpSqrt |
| --- | --- | --- | --- | --- |
| $10\sigma$ | mass | $1.52397 \times 10^{-23}$ | $7.21354 \times 10^{-7}$ | $0.000326548$ |
| | variance | $1.55416 \times 10^{-21}$ | $0.0000830583$ | $0.0513746$ |
| $25\sigma$ | mass | $6.11339 \times 10^{-138}$ | $4.41948 \times 10^{-16}$ | $1.14081 \times 10^{-6}$ |
| | variance | $3.83308 \times 10^{-135}$ | $2.92285 \times 10^{-13}$ | $0.000934673$ |
| $50\sigma$ | mass | 0. | $1.95318 \times 10^{-31}$ | $1.67283 \times 10^{-9}$ |
| | variance | 0. | $5.02302 \times 10^{-28}$ | $5.03447 \times 10^{-6}$ |

### Wave speed with infinite variance

In Fig. 1, we compared dispersal distributions that were scaled to have the same variance. This approach fails for distributions such as the Cauchy, that fall away so slowly that they have infinite variance. However, we can find the Cauchy distribution that yields the same wave speed as a Gaussian with (say) *σ* = 2 (Fig. 3). On an infinite range, a Cauchy distribution leads to a wave with ever-increasing speed. However, the Cauchy distribution must necessarily be truncated for numerical calculations, and for any given truncation point, the wave does settle to a steady speed, which can be matched to that of a Gaussian. Figure 3 plots the standard deviation of the truncated Cauchy against *a=* 1 - 2 p, for truncation at 10*σ* … 100*σ* (black … red dots). For pushed waves (a < 2), fat tails slow down the wave, because long-distance migrants are lost, and so the truncated Cauchy must have a higher variance to maintain the same speed (a < 2, left of Fig. 3). For a pulled wave, however, a truncated Cauchy with a very small variance can maintain high speed, provided that it is truncated sufficiently far out. In the most extreme case shown here (truncation at 100*σ*, *α*=4; bottom right of Fig. 3), 0nly a fraction 7.9 * 10"^8^ disperse away from their native deme, contributing variance ~10--^5^, and yet this maintains a wave with the same speed as Gaussian dispersal with standard deviation σ=2. (Note that for these extreme cases, the choice of deme spacing does affect the results, though not qualitatively).

### Random fluctuations

With long-tailed dispersal, deterministic models are misleading, because the numbers of long-range migrants will be small even in very dense populations. Indeed, even with Gaussian dispersal, finite population size has an appreciable effect on the speed of a pulled wave, because the dynamics depend on regions where alleles or individuals are very rare. Here, we simulate a finite population on an infinite range. The population is represented as a list of the numbers of the focal allele in each deme; all demes to the left are assumed to be fixed, and all to the right are assumed to be at zero. In each generation, and in each deme, there is selection followed by random sampling of 2N genes, following the Wright-Fisher model. Dispersal is implemented as follows. For each of the *n_p_* demes that are currently polymorphic, the 2N genes are drawn from a parent population according to the dispersal function; the continuous dispersal distribution is rounded to the nearest integer. (For σ=2, as used here, this causes negligible error). These genes can be from arbitrarily far away. The same procedure is followed for 1000 demes to the left of the current set, and 1000 demes to the right. Finally, the list is trimmed to remove fixed demes on the left and on the right (but tracking the location of the leftmost polymorphic deme, which will tend to move to the right as the wave advances. This scheme is extremely efficient, since the random distances moved can be taken as a single draw of 2Nn_p_ random numbers from the dispersal distribution. The only approximations are that individuals live on a discrete grid, and that only ±1000 demes are followed on either side.

Table 2 summarises the effects of finite population size. For a pushed wave (a=0.5), random drift has little effect, unless deme size is very small (2 *N* = 5). For a pulled wave, there is a substantial slowing, even for demes of 2 *N* = 1000 individuals. Note that this stochastic effect means that long range dispersal no longer increases the speed of advance, even for the exponential square root. There is no sign that rare propagules move far ahead of the initial wave, producing the kind of stochastic advance that is expected for very fat-tailed distribution. The key conclusion is that random fluctuations have very little effect on the speed of “pushed“ waves. The slowing effect is greater when waves are “pulled“, and there is long-range dispersal. However, even then the effect is not as dramatic as suggested by the asymptotic theory.

**Table 2.** The wave speed in a finite population. This is estimated as the mean speed over 2000 generations. The last row shows the results for the deterministic model (Fig. 1) with k_max_ = 20 for the Gaussian and Laplace distributions, and *k*_m_ = 200forthe exponential square root.

| $\alpha = 0.5$ | $2N$ | Gaussian | Laplace | ExpSqrt | $\alpha = 4$ | Gaussian | Laplace | ExpSqrt |
| --- | --- | --- | --- | --- | --- | --- | --- | --- |
|  | 10 | 0.263 | 0.274 | 0.267 |  | 0.533 | 0.540 | 0.546 |
|  | 100 | 0.259 | 0.257 | 0.243 |  | 0.697 | 0.672 | 0.705 |
|  | 1000 |  |  |  |  | 0.716 | 0.704 | 0.746 |
| | ∞ | 0.255 | 0.250 | 0.230 | | 0.751 | 0.765 | 0.920 |

**References not cited in the main text.**

Brunet E, Derrida B, Mueller AH, Munier S (2006) Phenomenological theory giving the full statistics of the position of fluctuating pulled fronts. Phys. Rev. E 73: 056126.

Fisher, RA (1937) The wave of advance of advantageous genes. Annals of Eugenics 7: 355-369.

Stokes AN (1976) On two types of moving front in quasilinear diffusion. Mathematical Biosciences 31: 307-315.

## Appendix B Establishing a wave

In a bistable system, a wave can be established if the initial distribution is above some threshold. Alternatively, one might continuously release within some region: again, there will be some threshold required for establishment. If the intensity of the source is above this threshold, then the wave can be established even if the source is turned off at a finite time. A key comparison is the total number that need to be introduced to establish the wave, or (more simply) how long the source needs to continue to send out the same number as would be needed with an initial brief pulse. Our intuition is that an initial pulse would always be more efficient, since any release in regions below the threshold will decay over time: the key is to raise a sufficiently large region above the threshold so that it can contribute to further increase.

We represent an influx by a term *mf*[*x*] *q*, which represents migration from a deme fixed for *p* = 1: in each generation, a (scaled) fraction *mf(X)* is replaced by migrants. This ensures that allele frequencies do not rise above&& 1. If one follows population density, scaled relative to carrying capacity, and with an Allee effect that causes bistability, then one would add a term *λf*[*x*], and density could rise above carrying capacity. Here we assume a source *λf(X)q*, corresponding to replacement of a fraction of the population. This is appropriate if population size is regulated to a constant value after the release.

### Island model

Take the simple case of a single population, with an influx that replaces a fraction m of the population every generation. Let 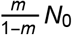

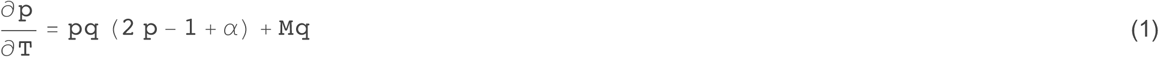

Then, it is easy to show that the critical migration rate is M^**\***^ = (1 - *α*)^2^/16. How much higher does *M* need to be if sustained for a finite time, *T*_o_? Integrating Eq. B1, starting from zero allele freequency:

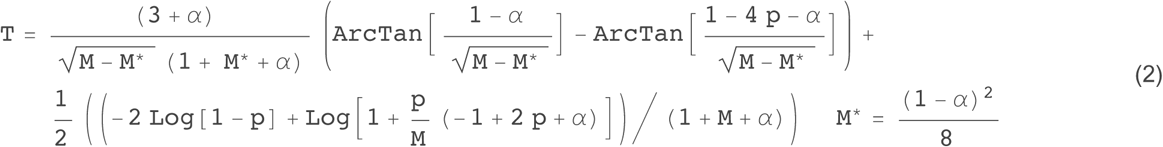

Now, establishment is assured if, at time *T*_crit_, 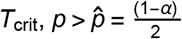. setiing 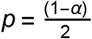

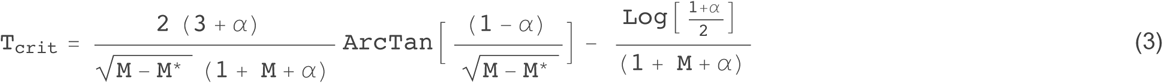

The left plot of Fig. B1 shows this critical time, as a function of the scaled migration rate, *M*. One might measure the total ^**‘**^cost^**’**^ of the introduction by *MT*. This declines to an asymptote for large M, indicating that the most efficient strategy is a short pulse that quickly raises the frequency above the critical threshold. However, *MT* is almost constant above *M*~0.1, and so the precise timing of the introduction makes little difference (Fig. B1, middle). If the resident population is constant, *N_0_*, then the number that need to be introduced to replace a fraction *m* is 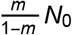. With complete CI, *s_h_ =* 1, and so,this equals 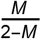: it becomes expensive to replace a large fraction of the population. When this effect is included (Fig. B1, right), the most efficient strategy is at intermediate *M*. However, an instantaneous pulse is almost as efficient. The best guidance is that the introduction should be made as rapidly as is feasible, given practical constraints.

**Figire B1:**
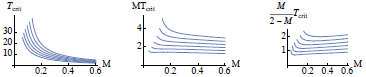
The left plot shows the minimum time needed for establishment, plotted against 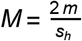, as a function of α=0.1,0.2, 0.3, 0.4, 0.5, 0.6 (bottom to top) (i.e., *p^̂^ =* 0.45, 0.4, 0.35, 0.3, 0.25, 0.2). The middle plot shows *MT*_crit_ *=* mtcrit, and the right plot 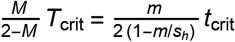

### One dimension

We rescale Eqs. 2, 3 by setting time to *T = (s_h_*/2) *t* and distance to 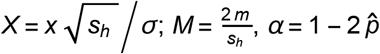

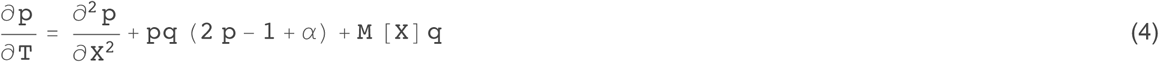

With M=0, the wave will increase if the initial p[x, 0] is “large enough” There is a critical bubble that lies on this theshold, which can be calculated explicitly in one dimension. Now, suppose that initially *p* = 0. How large must *M* be to ensure spread? This can be solved for special choices of the source, *M*[X]; below, we give the critical value for a point source, and for a “top hat” function, with a constant input within some region. We then give numerical results for the minimum time needed to ensure establishment, analogous to Fig. B1.

### Point source

With a source *Mδ(X)q,* there is a boundary condition 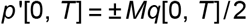. Integrating Eq. 1 we have, at *X* = 0:

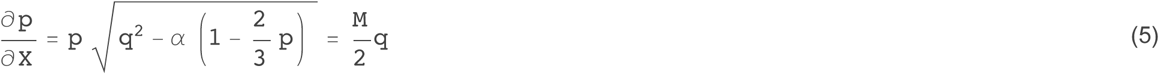

The function 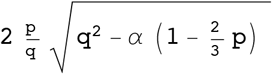 has an intermediate maximum, which gives the threshold value of Λ. Below this threshold, there are two solutions for p, one stable and one unstable:

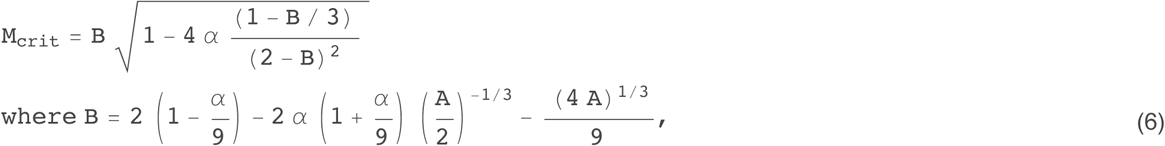

In the original variables, the threshold scales as 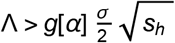 which has the dimensions *XT*^−1^

### Top-hat source

Now, suppose that we have a source *Mq* within *-Y*<*X*< *Y*. Then, equilibrium is given by:

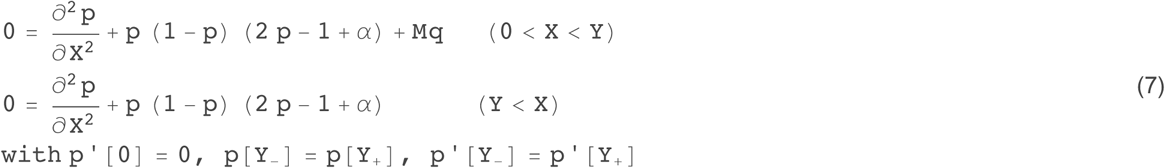

Intergrating:

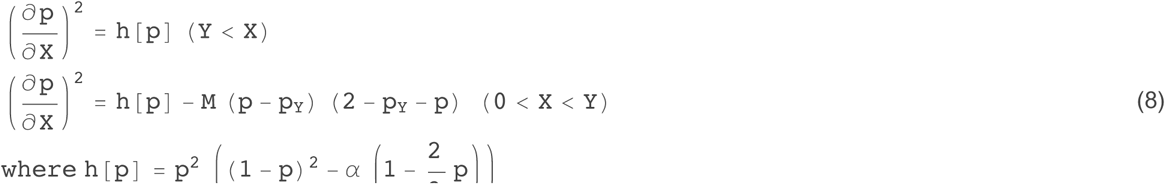

The allele frequency at zero, *p*_0_, iswhere*∂p*/*∂X* - 0, so that 0 = h[p_0_] - M (p_0_ - p_Y_) (2 - p_0_ - p_Y_); this defines *p_0_* as *a* function of *p*_Y_. We can obtain *Y* by integrating *dXIdp*. Hence, *p_Y_* is given by:

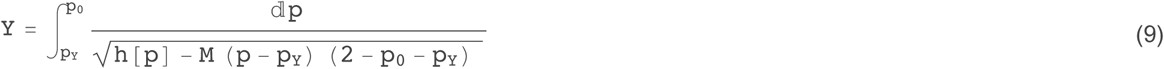

For given *p_Y_,* there is a solution for *Y,* provided that 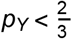; this has a maximum value of Y_cr_it, so that if *Y*< Y_crit_, there will be two solutions. This maximum defines the threshold for spread. However, it is not straightforward to find this threshold analytically.

Consider the equation 0 = h [p_0_] = M (p_0_ = p_Y_) (2 = p_0_ = p_Y_). For *M>* (1 = a)^2^/8, there is only one root. For *M<* (1 = a)^2^/8, there is one root if *p_Y_ >* p*_Y_.

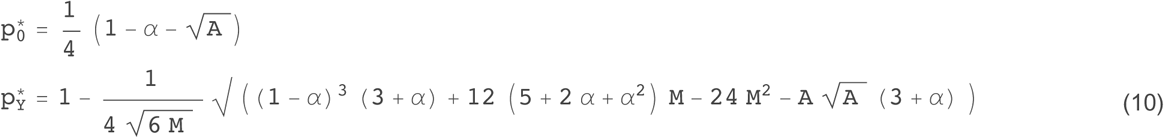

where A = (1 -α)^2^ - 8 M.

For *M>* (1 - α)^2^/8 = 0.02, there is a single solution, with a peak which represents the maximum Y_crit_ consistent with a stable equilibrium. However, once M passes below the threshold, there is a singularity, and it appears that there can be a stable solution with extremely large Y. This threshold simply corresponds to *M* so low that it cannot cause a transition, even as *Y*→∞ ro. Fig. B2 shows solutions to Eq. 10 over a range of *a, p;* the maximum gives the critical spatial range, Y_crit_, required for establishment. Table B1 gives this maximum, together with the total input, 2 MY_crit_. For given *a, p*, there is an optimal migration rate and spatial extent that minimises 2 MY_crit_. These analytic results give the minimum input required for establishment. However, it would take an indefinitely long time for establishment to be assured. If the input is more intense and/or over a wider spatial range, then it could persist for a shorter time. Results for a finite time, analogous to the island model (Fig. B1), require numerical solution. We give those in the next section, for two dimensions.

**Figure B2.**
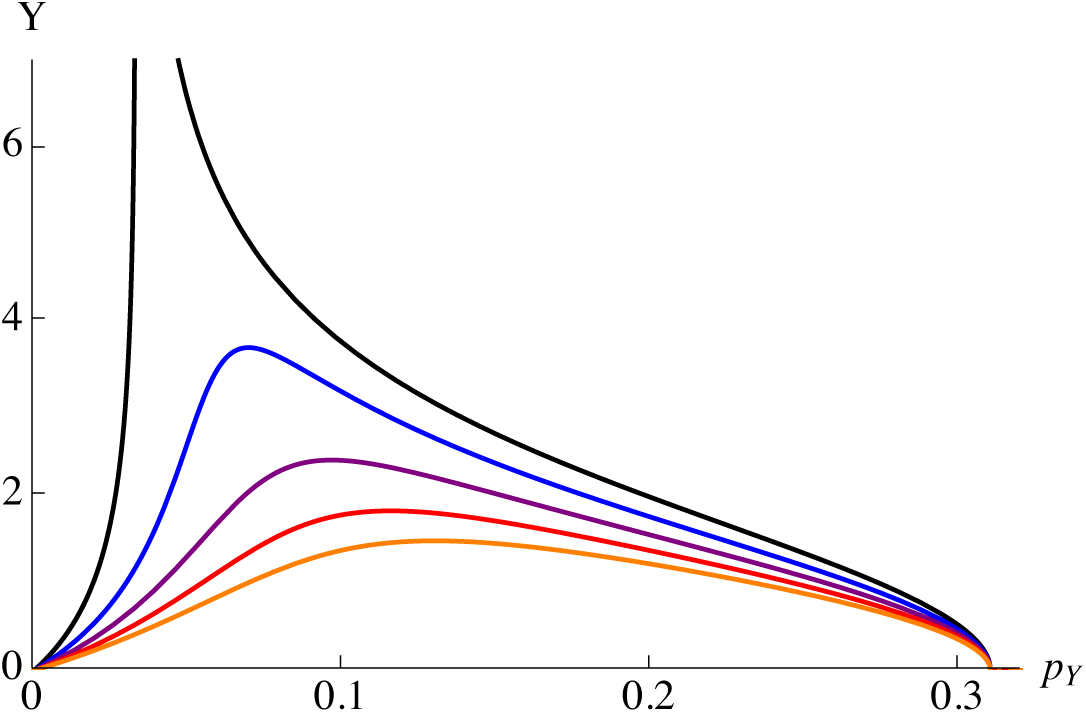
The value of Y against *p*_Y_, for α=0.2, p=0.2, and *M*= 0.02, 0.03, 0.04, 0.05, 0.06 (top to bottom). The maximum *Y* corresponds to the minimum size of the range, {-Y, Y} necessary for establishment, given a source M.

**Table B1.**
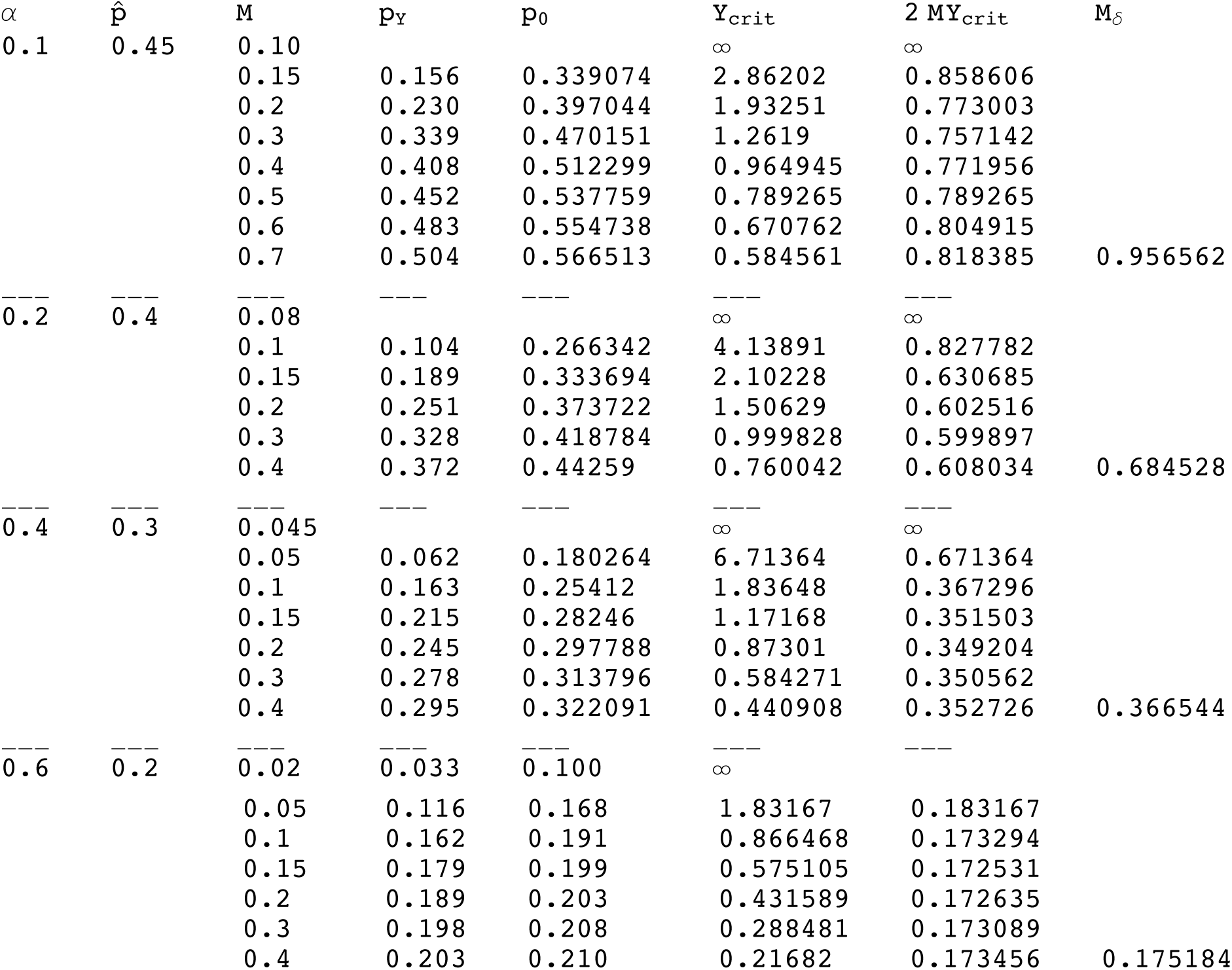
Threshold Y_crit_ for varying *a,* M. *p_Y_, p_0_* give the frequency at he edge of the source and at the centre, respectively. 2 *MY* gives the total rate of input. The last column, *M_6_,* gives the critical intensity of point source needed to allow establishment; it is the limit of the the total source 2 *MY* when *M* becomes large and its spatial extent, {-Y, Y} small.

### Two dimensions

Using the same scalings as in one dimensions, we have:

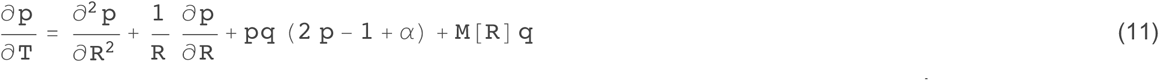

We solve this equation numerically using NDSolve in *Mathematics*. For a given *a, p^̂^,* there is a critical radius, R_crit_, such that the wave will just establish if initially *p* = 1 within *R <* R_crit_. We compare this initial pulse with a continuous source within the same radius, and sustained for time *T*; for given *T*, we find the minimum *M* required for establishment; this also gives the minimum T_crit_ needed for establishment with given M. As *M* increases above the threshold required for establishment from an indefinitely sustained source, T_crit_ decreases. Figure B3 summarises these results, in the same form as for Fig. B1, but just for *a* = 0.6, p ^*̂*^= 0.2. As for the island model, the effort, measured by *MT* reaches an asymptote for large M, indicating that an immediate increase to high frequency is most efficient. If the increasing cost of raising frequency by introduction into a fixed native population is included, by using he measure 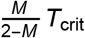, then the most efficient scheme is to use *M* ~ 0.3 for T_crit_ ~ 1 - which is still an extremely short time if *s*_***h***_*=* 1.

**Figure B3.**
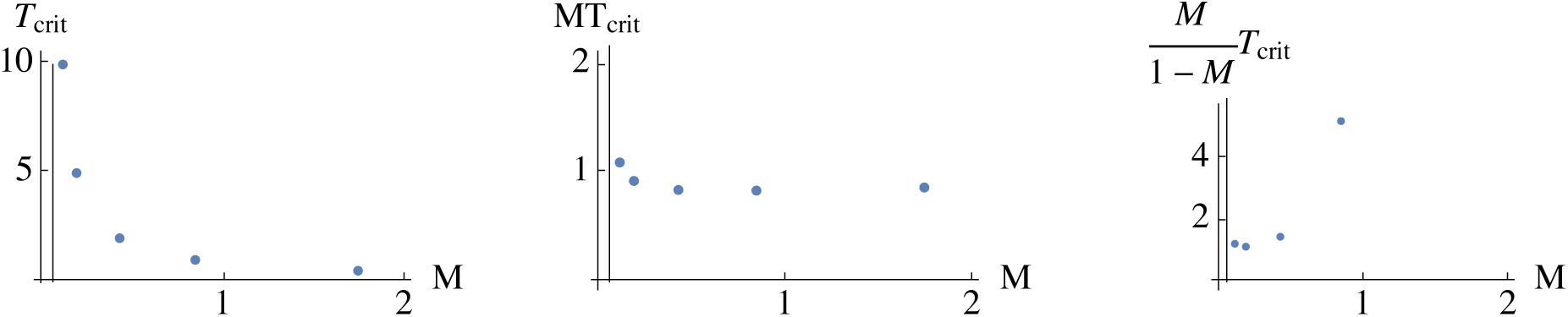
On the left, the time for which a source needs to be sustained, plotted against source strength, M, given optimal initial radius. The vertical line is the source strength needed for establishment, if sustained indefinitely. The middle plot shows the total amount, 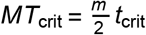 that is needed, as a function of source strength. The right plot shows 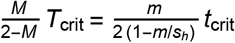, which is the total number that need to be introduced, relative to a constant native population, if *s*_***h***_*=* 1. R_crit_ = 5.19, *a* = 0.6, *p^̂^ =* 0.2.

